# A spatial atlas of chemoradiation therapy in pancreatic cancer identifies cellular and microenvironmental determinants of persister populations

**DOI:** 10.1101/2025.06.20.660757

**Authors:** Vincent Bernard, Li-Ting Ku, Tianyu Wang, Ariana Acevedo-Diaz, Kimal I Rajapakshe, Galia Jacobson, Daniela Tovar, Jimin Min, Guangsheng Pei, Candise Tat, Ayush Suresh, Ching-Wei D. Tzeng, Matthew HG Katz, Manoop S. Bhutani, Huaming Wang, Robert A Wolff, Cara Haymaker, Ethan B Ludmir, Huocong Huang, Xiongfeng Chen, Liang Li, Albert C. Koong, Linghua Wang, Nicholas E Navin, Dadi Jiang, Ziyi Li, Anirban Maitra, Eugene J Koay

## Abstract

The molecular pathways involved in the response to radiation therapy in pancreatic ductal adenocarcinoma (PDAC) remain poorly understood. We aimed to elucidate the adaptive mechanisms and cellular interactions within PDAC to radiation therapy (RT). We constructed a transcriptomic landscape of the cellular subtypes and spatially resolved neighborhoods from 50 patient samples, including 16 longitudinally matched single cell RNA sequencing and 34 spatial transcriptomics specimens. To resolve shortcomings of cell-type mixtures in spatial data, we developed a novel statistical method called SpaCCI (spatially aware analysis of cell-cell interactions) to profile cell–cell interactions and ligand-receptor enrichment. This revealed CXCL12/TGFβ-driven persister cell niches where activated fibroblasts reprogram tumor-associated macrophages and spatially exclude stress-response CD8 T cells after RT. Persister cancer cells displayed transcriptional evidence of recalcitrance to metal-induced cell death pathways of ferroptosis and cuproptosis which were recapitulated in preclinical models. Our study reveals the selective pressures experienced by PDAC following RT that may help provide insight for future multimodal therapeutic strategies.

## Introduction

Pancreatic ductal adenocarcinoma (PDAC) is among the most lethal malignancies, with a dismal 5-year survival rate^1^. This outcome is driven in part by PDAC’s intrinsic resistance to therapy as well as its uniquely dense and immunosuppressive tumor microenvironment (TME), which impedes effective delivery of therapies and fosters therapeutic resistance^2^. Radiotherapy (RT) continues to have a role in PDAC management, but several modern clinical trials suggest the efficacy of RT may be limited for patients with non-metastatic PDAC (e.g., confined to the pancreas)^3, 4^. Indeed, while PDAC tumors typically exhibit radio-resistance, the molecular pathways contributing to this phenomenon remain poorly understood despite extensive investigation^5-9^ Understanding the cell intrinsic and microenvironmental pathways that orchestrate the relative recalcitrance of PDAC to RT could provide unprecedented insights into the design of combination regimens. Here, we apply high-resolution single-cell and spatial transcriptomics profiling in a series of PDAC samples (including a subset of matched samples before and after RT), aiming to delineate the TME alterations and resistant intrinsic cell states that emerge after radiotherapy. In particular, we investigate so-called *persister* cell populations, the critical subfraction of cancer cells that survive RT, characterizing their adaptive molecular mechanisms and complex interactions within the TME that contribute to therapeutic evasion. To facilitate our assessments, we also leverage a novel analytical platform called Spatially Aware Cell-Cell Interaction (SpaCCI), which integrates spatially-resolved transcriptomics with computational cell– cell interaction mapping to characterize ligand–receptor signaling networks in situ within the TME. Using SpaCCI, we profiled the localization and communication of immune, stromal, and tumor cells, enabling us to detail critical cell–cell interactions and signaling pathways associated with therapy resistance. Together, this study represents the first comprehensive single cell and spatial atlas of PDAC following neoadjuvant RT (including a range of dosing and RT modalities), which serves as the seedbed for approaches that could improve treatment responses, and potentially survival in patients.

## Results

### Single cell and spatial transcriptomics from human PDAC tumors

We collected 16 serial fresh primary PDAC tissues from 9 patients (7 with matched samples) from a previously published phase Ib/II trial of patients who underwent neoadjuvant therapy with 12 weeks of chemotherapy followed by stereotactic body radiation therapy (SBRT) with 50Gy in 5 fractions (Figure 1A)^10^. The first timepoint for single cell RNA sequencing (scRNA-seq) from a core biopsy was after neoadjuvant chemotherapy, but before SBRT. The second timepoint for scRNA-seq was approximately 12 weeks after the end of SBRT. Additionally, we performed spatial transcriptomics on 34 PDAC formalin fixed paraffin embedded tissues taken at the time of primary tumor resection for patient that either received upfront surgery and thus received no neoadjuvant treatment (n=11), or received neoadjuvant therapy including chemoRT with 30Gy in 10 fractions (n=7), 50Gy in 25 fractions (n=9), or SBRT (40-50Gy in 5 fractions) (n=7) (**Figure 1A**).

**Figure 1:**
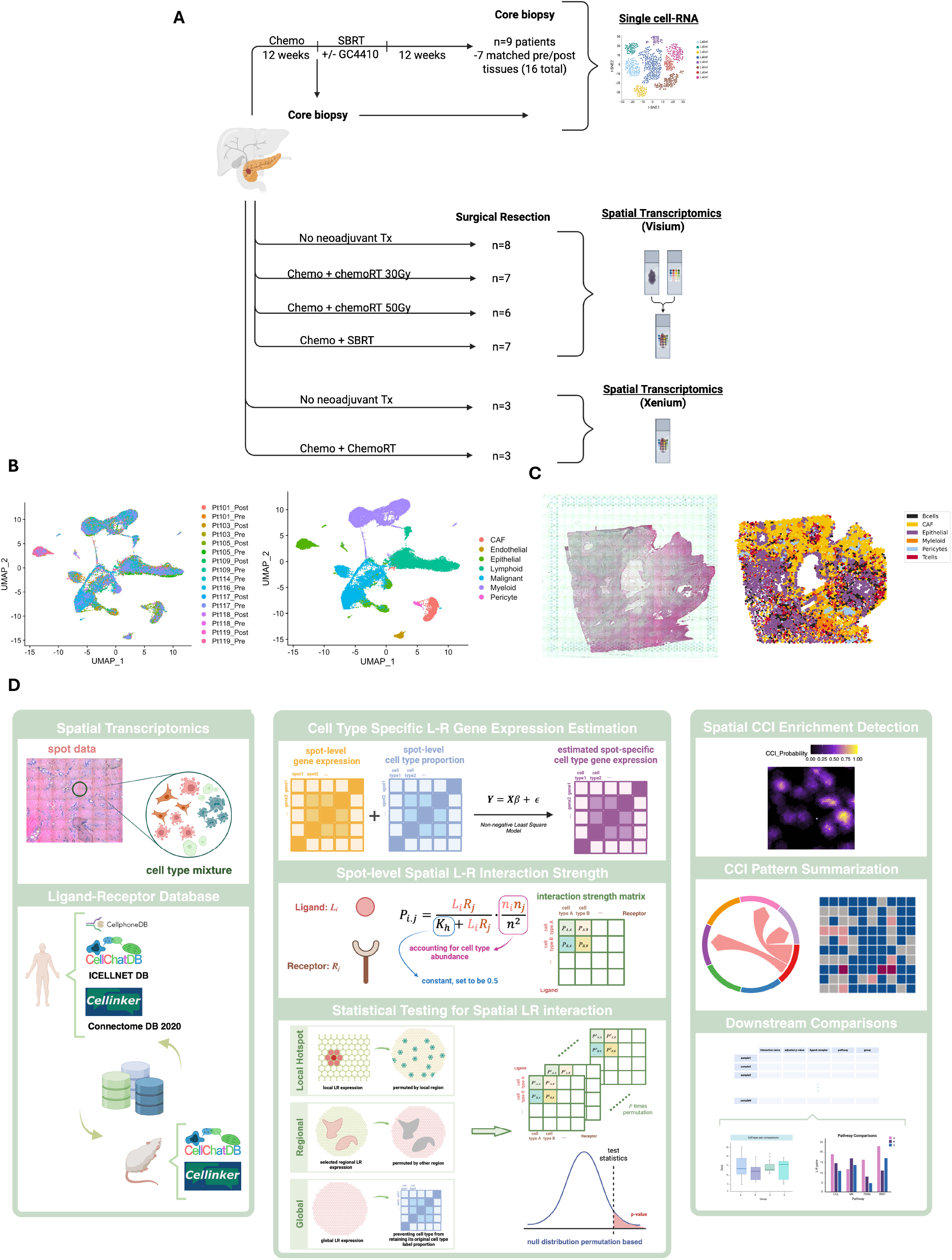
A. Schematic overview of human tissue samples taken to develop single cell and spatial transcriptomic datasets. B. UMAP of scRNA depicting distribution of patients (left) and cell types (right). C. Cell types as identified by scRNA signatures were used for deconvolution of cells in spatial transcriptomic dataset. D. Schematic workflow overview of Spatially Aware Cell-Cell Interaction (SpaCCI) pipeline.

Unsupervised clustering of our scRNA-seq dataset revealed 7 main clusters of cells representing neoplastic cells and non-malignant cell types including, cancer associated fibroblasts (CAFs), endothelial cells, epithelial cells, lymphoid, myeloid cells, and pericytes, (**Figure 1B**). Inferred copy number variation (inferCNV) per an estimated copy number score confirmed neoplastic cell type clusters^11^. We further cataloged cells following subset analyses based on differential gene expression profiles leading to a total of 45 unique cell types. We then performed deconvolution of the derived signatures from these cell types on the spatial transcriptomics dataset utilizing three independent algorithms: Cytospace, RCTD, and iSTAR^12-14^(**Figure 1C**). This allowed us to profile correlations of cell types with treated or untreated PDAC tissues (**Figure 2A**). Specifically, if cell types achieved a odds ratio (OR) cutoff of >1.25 or <0.75 and p-value of <0.05, and were concordant in both single cell and spatial datasets, these were determined to be of significance for further profiling as cells that were significantly enriched or depleted following RT.

**Figure 2:**
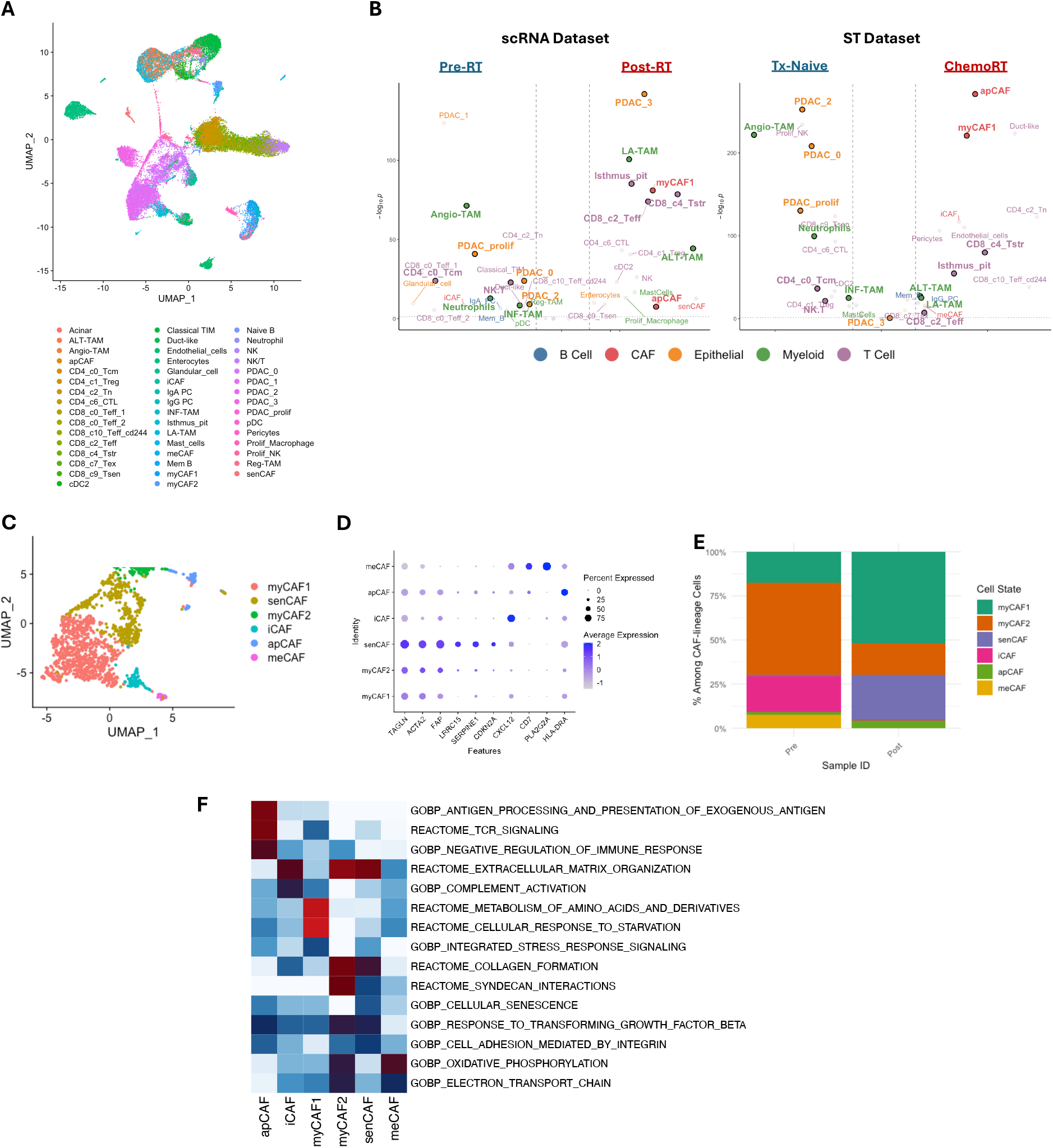
A. Annotated cell types identified in the single cell dataset used for deconvolution. B. Volcano plot of odds ratio (OR) for cell types in the context of pre and post radiation therapy in scRNA dataset or treatment naïve vs chemoRT in spatial transcriptomic dataset. Grey dashed lines mark the enrichment depletion thresholds (OR < 0.75 or > 1.25 and pvalue < 0.05). Cell types with common patterns across both datasets are accentuated. C. UMAP of subsetted CAFs in scRNA dataset with respective D. markers genes. E. Bar graph demonstrates kinetics of CAF proportion before and after radiation therapy. F. Heatmap of significantly enriched pathways characterizing function of detailed CAF subtypes

### Radiotherapy Reshapes CAF Subpopulations and Stromal Signaling in PDAC

Within the scRNA-seq dataset we identified six unique populations of CAFs characterized by distinct gene expression patterns (**Figure 2B-D**). In PDAC tissues receiving RT, the CAF compartment shifts toward a higher proportion of antigen-presenting CAFs (apCAFs) (OR 1.84, CI 1.78 – 1.91, pval <0.0001) and myofibroblastic CAF1 (myCAF1) (OR 1.74, CI 1.69 – 1.81, pval <0.0001) cells(**Figure 2B,E**). Deconvolution of CAF population signatures in our spatial transcriptomic dataset independently validated these compartment shifts. The apCAF subset shows elevated expression of MHC class II antigen presentation genes (HLA-DRA) and is enriched in pathways for exogenous antigen processing and T-cell receptor signaling (**Figure 2F)**^15, 16^. Concurrently, myCAF1 cells display a prominent myofibroblastic program, marked by high α-smooth muscle actin (*ACTA2*), abundant extracellular matrix components (*COL1A1, COL1A2*, and *COL3A1*), and contractile proteins (*TAGLN, MYL9*), consistent with a contractile, matrix-depositing phenotype. These RT-altered CAF populations also activate nutrient stress-response pathways (such as GCN2–EIF2α signaling), reflecting gene programs for amino acid deficiency and starvation responses^17^. Notably, senCAFs characterized by increased *LRRC15, SERPINE1*, and *CDKN2A* expression remain prevalent or increase after RT, with enrichment in TGFβ-response and integrin-adhesion pathways (**Figure 2B, E, F**)^18^. These features of post-RT senCAFs are consistent with reports that senescent myofibroblasts accumulate during PDAC progression and drive fibrosis and immunosuppression in the TME.

### Radiation induces an expanded stress response state among tissue-infiltrating T cells

scRNA-seq profiling of matched PDAC tumors before and after RT revealed shifts in intratumoral T-cell composition classified based on previously reported signatures of tumor infiltrating T cells (**Figure 3A**) ^19^. In terms of CD8+ T-cell populations, the relative frequencies of stress response (CD8_c4_Tstr) (OR 1.97, CI 1.83 – 2.11, pval <0.0001) and effector CD8+ T cells (CD8_c2_Teff) (OR 1.32, CI 1.20 – 1.47, pval <0.0001) were significantly increased **Figure 2B, 3B)**. The increase in CD8_c4_Tstr population following RT was notable for heightened stress-response, chemokine receptor signaling, moderate cytotoxicity, and activation-effector function signal (**Figure 3C, D**). This subset displayed elevated heat-shock proteins HSPA1A/HSPA1B and chronic stimulation markers (CD69, RGS1), consistent with a state of persistent activation. These Tstr cells upregulated proteotoxic stress pathways (chaperone-mediated protein folding and HSF1 activation). Notably, stress-response CD8 T-cell populations have been linked to aggressive tumor subtypes and poor response to immune checkpoint blockade^19^. CD8 subsets Teffector populations show strong expression of cytotoxicity and effector activation. The CD8_c2_Teff cluster exhibited a robust cytotoxic phenotype, marked by high expression of granzyme B (GZMB), perforin (PRF1), and granulysin (GNLY) ^20^. Collectively, these features reflect potent cytotoxic activity and chemotactic signaling that promote anti-tumor immune responses in PDAC. However, spatial transcriptomics revealed exclusion of effector T cells from the tumor bed, suggesting mechanisms of active immune exclusion. (**Figure 3E**). NK/T and NK cells displayed effector and cytotoxic potential with moderate metabolic activity, suggestive of anti-tumor activity, while Prolif_NK cells demonstrated greater metabolic activity and stress response (**Figure 3C, D**).

**Figure 3:**
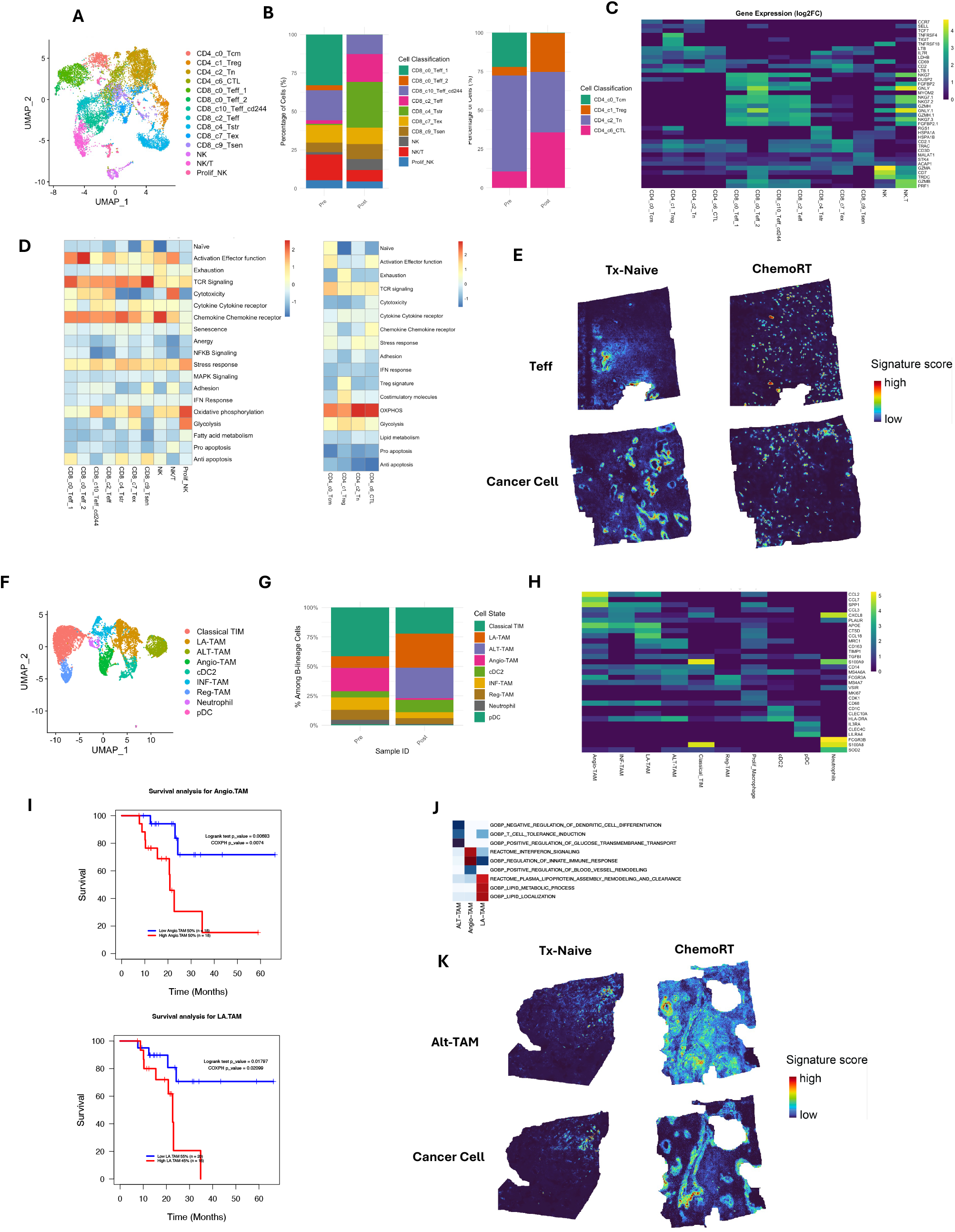
A. UMAP of subsetted T & NK cells in scRNA dataset B. Bar graph demonstrates kinetics of cell proportions before and after SBRT among CD8 (left) and CD4 (right) subsets. C. Heatmap of differentially expressed genes amongst all T-cell subtypes. D. Heat map illustrating expression of curated gene signatures across CD8^+^ (Left) and CD4^+^ T cell clusters. Heat map was generated based on module scores. E. Spatial visualization of malignant cells and T cell signature scores imputed by inferring Super-resolution Tissue ARchitecture (iSTAR) from representative naïve and chemoRT treated sample demonstrating exclusion of T effector populations from tumor bed. F. UMAP of subsetted myeloid cells in scRNA dataset. G. Bar graph demonstrates kinetics of cell proportions before and after SBRT of selected myeloid populations. H. Heatmap of differentially expressed genes amongst all myeloid subtypes. I. Kaplan meier curve of TCGA dataset of patients treated with chemoRT demonstrating decreased overall survival with greater proportion of Angio-TAM (logrank pval <0.01) and LA-TAM (logrank oval =0.02) signatures in resected tumors. J. Heatmap of significantly enriched pathways characterizing function of selected myeloid subtypes. K. Spatially representation of ALT-TAMs demonstrating enrichment within the tumor bed following chemoRT when compared to treatment naïve.

Central memory (CD4_c0_Tcm) populations exhibit elevated TCR signaling signatures alongside expression of central memory-associated markers such as CCR7, SELL (CD62L), and TCF7, indicative of a long-lived, lymph node-homing, memory T-cell phenotype that contracts following RT (OR 0.66, CI 0.62 – 0.70, pval <0.0001). Regulatory CD4 (CD4_c1_Treg) T cells distinctly show heightened expression of Treg-related signatures, as well as OXPHOS and glycolysis, suggesting metabolic flexibility linked to immune suppression (**Figure 3D**). Cytotoxic (CD4_c4_CTL) cells are characterized by effector function, receptor signaling and OXPHOS activity. Naïve-like (CD4_c2_Tn) cells show relatively low activation and cytotoxicity.

### Radiation Induces TAM Subtype Redistribution and Immune-Evasive Signaling in PDAC

Post-RT tumors showed a notable decrease in infiltrating neutrophils, alongside modulated macrophage populations (**Figure 2B, 3F-G**). Importantly, neutrophils have previously been demonstrated to promote resistance to RT with lower absolute neutrophil counts associating with better patient outcomes in solid tumors^21, 22^. Tumor-associated macrophage (TAM) subpopulations were redistributed following RT. The pro-angiogenic Angio-TAM subset was significantly diminished, whereas alternatively activated (ALT-TAM) and lipid-associated (LA- TAM) subsets became more prevalent (**Figure 2B, 3G**). Notably, patients whose tumors were enriched in Angio-TAMs (SPP1^+^ macrophages) and LA-TAMs (APOE^+^ macrophages) after radiotherapy had shorter survival outcomes (**Figure 3I)**.

Angio-TAMs secrete high levels of chemotactic and pro-angiogenic factors (CCL2, CCL7, SPP1), which drive monocyte recruitment and neovascularization in the tumor (**Figure 3H**)^23^. By contrast, LA-TAMs are characterized by elevated apolipoproteins and lysosomal enzymes (APOE, CTSD) together with immunosuppressive mediators such as CCL18. Pathway analysis indicates that LA-TAMs are associated with lipid metabolism and vesicle transport pathways (plasma lipoprotein remodeling, receptor-mediated endocytosis) (**Figure 3J**). ALT-TAMs upregulate immunomodulatory cytokines and extracellular matrix remodelers including TGF-β, MRC1 (CD206), CD163, and TIMP1 (**Figure 3H**). Consistent with this phenotype, ALT-TAMs are enriched in immunoregulatory and metabolic pathways (negative regulation of dendritic cell differentiation, induction of T-cell tolerance, and positive regulation of glucose transport) (**Figure 3J**), supporting tumor progression and fibrotic stroma formation. The increased prevalence of ALT-TAMs directly juxtaposed to tumor cells following RT suggests that these immunosuppressive interactions occur immediately within the tumor bed **(Figure 3K)**.^24^.

### Cancer Cell Molecular Classification Reveals Persister Cell Populations with Metabolic Dependencies Following RT

ScRNA-seq analysis further identified five distinct malignant cell populations with differing post-RT kinetics: PDAC_0, PDAC_1, PDAC_2, PDAC_3, and PDAC_prolif (**Figure 4A, B**). Pathway analysis of all composite malignant cell population revealed enrichment of KRAS signaling, lipid metabolism, and adaptive immune response following RT (**Supplementary Figure 7A**). Cancer cells were further classified into “classical” or “basal” subtypes within both datasets based on computed signature scores derived from established PDAC molecular subtype signatures^25^. Tumor regions that exhibited overlapping markers, admixtures of cells from both subtypes, or low expression of subtype-specific markers were categorized as “intermediate”^26^. In post-RT tissues, basal-like subtypes were depleted (OR 0.24 P<0.0001), while intermediate and classical subtypes persisted in both datasets (**Figure 4C, Supplementary Figure 7B**). This was further demonstrated following deconvolution of classical and basal signatures through iSTAR demonstrates enrichment of basal subtypes in treatment naïve PDAC tissues, compared to a decrease in their prevalence in RT-treated samples **(Supplementary Figure 7C)**.

**Figure 4:**
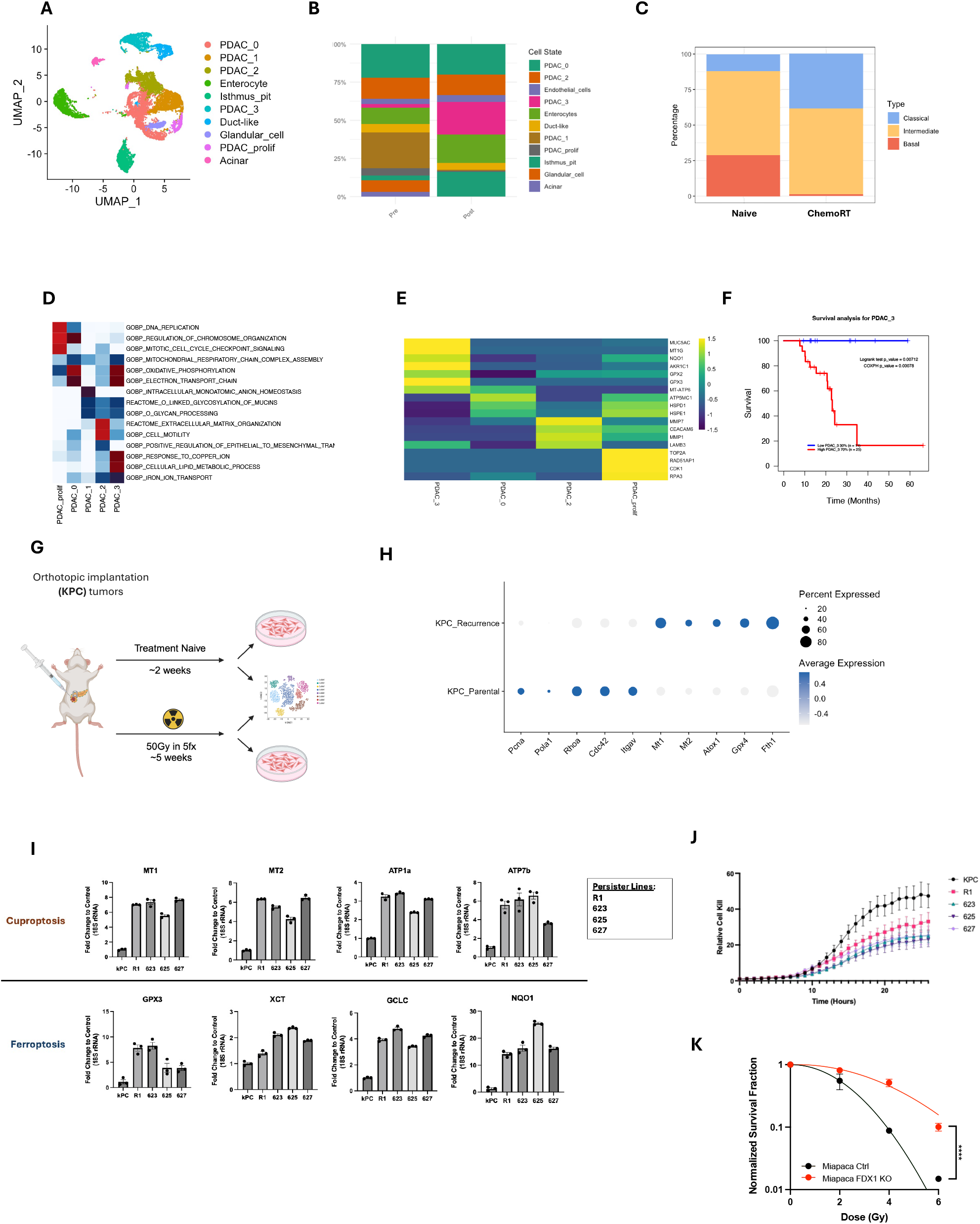
A. UMAP of single cell dataset demonstrating annotated epithelial cell types. B. Bar graph demonstrating changes in cell types as a proportion of all epithelial cells with a significant increase in PDAC_3 subtype of cancer cells likely representing a resistant cell type. C. Bar graph of module score across all spatial transcriptomic tissues demonstrates that ChemoRT induces ablation of basal-like molecular subtypes vs expansion and persistence of classicaland intermediate subtypes, respectively. D. Dot plot identifying significant pathways enriched in malignant cell types, particularly with PDAC_3 demonstrating enrichment of lipid and ion metabolism related pathways. E. Heatmap of cancer cell subtypes demonstrates highly expressed genes related to lipid metabolism, mitochondrial metabolism, EMT, and proliferation & DNA repair across cancer cell subtypes. F. Kaplan meier curve of TCGA dataset of patients treated with chemoRT demonstrating decreased overall survival with greater proportion of PDAC_3 subtype signature (logrank pval <0.001). G. Schematic representation of orthotopically injected KPC lines into syngeneic mice in order to develop persister cell lines (R1, 623, 625, and 627). H. Dotplot of gene expression from scRNA-seq from KPC parental and recurrent tumors demonstrating expression of representative gene programs of cell proliferation, curproptosis defense, and ferroptosis defence. I. Barplots demonstrating fold change relative to control (18S rRNA) of annotated genes in untreated (KPC) and persister cell lines. J. Relative cell kill as measured by Cytotox Dye following treatment with Elesclomol (15nM) in Copper (2uM) enriched media as measured by Incucyte. K. Clonogenic assay of FDX1 Knockout compared to parental Miapaca PDAC cell line.

Amongst the five malignant PDAC molecular subpopulations, PDAC_3 was shown to persist in both the scRNA-seq and spatial transcriptomic datasets following RT suggestive of a therapy resistant cancer subtype (**Figure 2B, 4B**). These cells exhibited gene expression signatures associated with lipid metabolism and copper ion homeostasis (**Figure 4D**). Key upregulated genes included MT1G and ATP7A, which play roles in copper detoxification, alongside NQO1, GPX2, and GPX3, which provide defense against oxidative stress (**Figure 4E**)^27^. Given that these genes and pathways represent protective mechanisms against iron and copper mediated cell death, these findings suggest that PDAC_3 cells may evade ferroptosis and cuproptosis through specific metabolic adaptations^28, 29^. Furthermore, deconvolution of the PDAC_3 signature in tissue samples from a subset of RT-treated patients within the TCGA dataset revealed that patients with a higher proportion of this signature post-RT experienced poorer survival outcomes (**Figure 4F**). In contrast, PDAC_prolif, characterized by enrichment in cell cycle and DNA replication pathways, and PDAC_2, associated with extracellular matrix (ECM) remodeling and epithelial-to-mesenchymal transition (EMT), were significantly reduced in post-RT samples (**Figure 2B, 4D**). PDAC_0, enriched for oxidative phosphorylation pathways, also exhibited reduced prevalence following RT.

### Persister Cell Model Demonstrates Cuproptosis and Ferroptosis Defense Pathways Following Radiation

To validate the PDAC_3 persister cell phenotype, we established orthotopic KPC tumors in C57BL/6 mice and treated them with a clinically relevant dose of 40Gy in 5 fractions of RT (Figure 4G). We harvested tumors at the time of recurrence following RT and performed scRNA seq as well as established in vitro cell lines. ScRNA-seq recapitulated increase in expression patterns of cuproptosis (Mt1, Mt2, and Atox1) and ferroptosis (Gpx4 and Fth1) defense genes in the recurrent tumors compared to the non-treated parental KPC tumors (**Figure 4H**). Quantitative RT-PCR of established cell lines demonstrated upregulation of cuproptosis defense genes (Atp7a, Atp7b, Mt2) and ferroptosis defense genes (Gpx3, Slc7a11-xCT, Nqo1, Gclc) in the RT-treated tumor lines, as compared to an untreated KPC cell line **(Figure 4I)**^27^. Western blots also demonstrated increased expression of ATP7A and ATP7B, key copper transport proteins, MT2, a known copper sequestering protein, and xCT (SLCA11) a ferroptosis defense membrane antiporter, indicating enhanced cuproptosis and ferroptosis resistance **(Supplementary Figure 7D)**. Treatment with the cuproptosis-inducing agent Elesclomol in copper-enriched media demonstrated reduced sensitivity of persister cell lines to cuproptosis-induced cell death compared to parental KPC cells **(Figure 4K)**. Knockout of FDX-1, a gene essential for cuproptosis induction^30^, in MIA PaCa-2 resulted in increased radioresistance based on colony formation assay, further suggesting an essential role for cuproptosis related cell death in response to radiation therapy **(Figure 4L)**.

### Development of a spatial cell-cell interaction platform

The Spatially Aware Cell-Cell Interaction (SpaCCI) pipeline is illustrated in Figure 1D. Please also see Methods and Supplementary Materials for more details. Briefly, SpaCCI is an integrative statistical model designed for quantifying cell-cell interactions by incorporating spot-level cell type-specific gene expression and spatial information. SpaCCI builds upon the principle that each spatial domain on the tissue comprises a mixture of cell types, with ligands and receptors interacting across cells within limited spatial ranges. This is unlike traditional methods that often analyze spot-level data similarly to single-cell transcriptomics. Consequently, SpaCCI integrates transcriptomics measurements with the cell type composition of each spatial domain and ligand-receptor databases to ensure accurate and interpretable detection of cell-cell interactions. Importantly, by retaining and utilizing spatial information from neighboring spots, SpaCCI can quantify both local and global interaction patterns, capturing the intricate interplay between cells that shapes biological processes.

To evaluate SpaCCI’s performance, we conducted simulations using previously published mouse brain Stereo-seq data^31^ and pancreatic cancer scRNA-seq data^32^. In the mouse brain dataset, SpaCCI effectively captured localized and global interaction patterns, aligning with known spatial organization and outperforming existing methods such as CellChat and CellPhoneDB in accuracy (**Supplementary Fig. 2**). SpaCCI robustly captured realistic cell-type compositions within simulated spot-level datasets derived from mouse brain Stereo-seq data, reflecting proportions and distributions observed in actual tissues (**Supplementary Fig. 3**). It accurately recovered ground-truth cell-cell interaction hotspots, demonstrating effective spatially-localized detection (**Supplementary Fig. 4**). We evaluated SpaCCI’s global detection capability by comparing its inferred cell–cell interaction patterns with those derived from CellChat and CellPhoneDB, using adult mouse brain Stereo-seq data. SpaCCI demonstrated strong consistency with both methods (correlations of 0.85 and 0.89, respectively), highlighting its robustness in identifying global interaction patterns across different ligand-receptor databases (**Supplementary Fig. 5**). These findings highlight SpaCCI’s robust ability to infer biologically relevant cell-cell interactions across diverse datasets and disease contexts.

### Distinct ligand-receptor signatures characterize spatial communication after RT in PDAC

We applied SpaCCI to each sample individually due to the distinct spatial contexts and dynamics present in each sample **(Supplementary Fig. 8 - 11)**. In profiling global ligand receptor changes across all cell types, after aggregating the results within each treatment group (Naïve vs ChemoRT: SBRT, CRT30 and CRT50), we observed a significant increase in cell-cell interactions following chemoRT (**Fig. 5E, 5F**). Specifically, the ChemoRT treatment group exhibited enriched interactions of myeloid cells with T cells, T cells with endothelial cells, and T cells with B cells, compared to the Naïve group (**Supplementary Fig.12A,B)**. This increase in cell-cell interaction counts was also confirmed in our scRNA dataset **(Supplementary Fig. 13A**). Further analysis of ligand-receptor interactions between the Naïve and chemoRT groups revealed significant enrichment of several interaction pathways which were also observed in our scRNA dataset **(Figure 5G, Supplementary Fig. 13B**). Specifically, we observed a surge in TGFβ interactions between T cells, B cells, and endothelium, which may have pro- or anti-tumorigenic mechanisms in the TME depending on context, as well as an enrichment of TGFβ family signaling involving CAFs after RT, consistent with myCAF-driven fibrosis and matrix production^33, 34^. Additional putative pro-tumoral axes such as MIF–CD74, IL16-CD4, and CXCL12–CXCR4 are active in the irradiated stroma, supporting immunologic stress and excluding cytotoxic T cells from tumors^35, 36^. More specifically, CD8_Tstr cells were seen to engage in TGFB1–TGFBR1, CXCL12–CXCR4, and MIF–CD74/CXCR4 interactions with CAFs and TAMs indicating adaptation to tumor-associated stress within the tumor microenvironment. SpaCCI then allows for visualization of co- localized interactions between predicted cell types. This demonstrates significant enriched cell type pairs after ChemoRT showing more spatially concentrated interaction signals compared to treatment-naïve tissues, highlighting the differential impact of treatment on cellular communication dynamics (**Figure 5H**).

**Figure 5:**
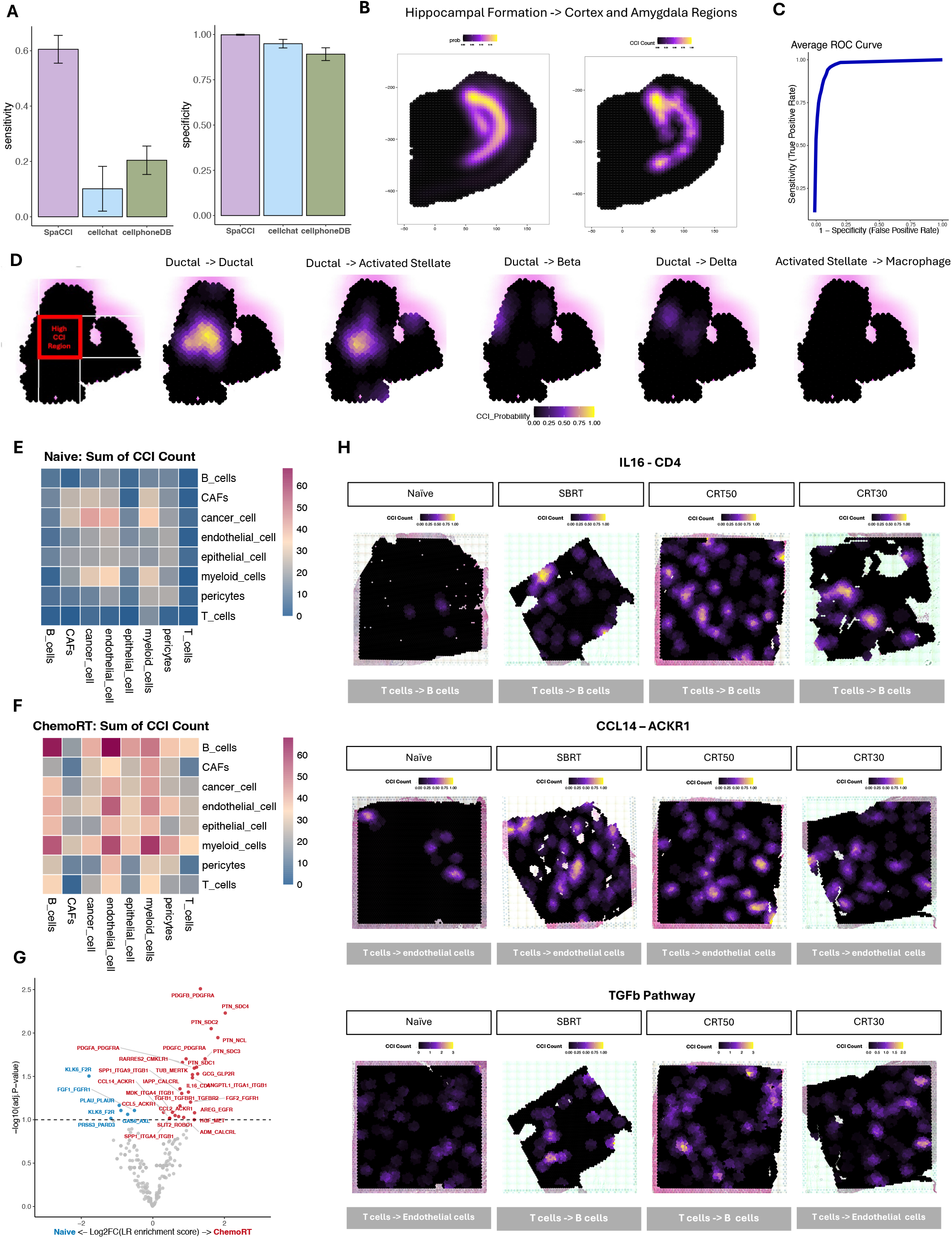
A. The sensitivity and the specificity comparisons among SpaCCI, cellchat and cellphoneDB LR pipelines with simulated spot dataset. B. The true interaction strength (left) from the pseudo stereo-seq data and the detected interaction strength (right) from SpaCCI, showing that SpaCCI captures interactions similar to what is expected. C. The average ROC curve of the localized detection using SpaCCI. D. A simulation demonstration was generated simulated data should have high CCI occurring with specific CCI directions (left). The simulation results from SpaCCI showing that as expected, we detect the directions of Ductal to Ductal and Ductal to Activated Stellate cell-cell interaction in the center region. E, F. Heatmap of aggregate results from SpaCCI in Naïve and ChemoRT samples respectively. G. Volcano plot of LR interaction pair comparisons between ChemoRT samples and Naïve samples. H Localized spatial plot of the IL16-CD4, CCL14-ACKR1, and TGFb pathway LR pair across treatment groups.

Focusing SpaCCI on cancer cell populations, PDAC_3 cells showed enrichment in pathways related to Sema3 in the chemoRT group (ER 1.87, pval = 0.03), when compared to the treatment naïve group (ER 1.3, pval = 0.43) **(Supplementary Figure 14A**). This was specifically related to Semaphorin 3 ligand activity (SEMA3C) engaging Neuropilin-1/2 and PlexinA1-1 receptors pathways known to modulate angiogenesis, immune responses, and therapy resistance (**Supplementary Figure 14B)** ^37, 38^. These interactions were confirmed in our scRNA-seq dataset where significant enrichment in Sema3 pathways were observed in post-RT tissues (ER 2.28, pval <0.01) compared to pre-RT tissues (ER 1.61, pval=0.14) (**Supplementary Figure 14C, Supplementary Fig. 13C**). An increase in overall Sema3 interactions across all cell types were observed following RT (**Supplementary Figure 14D,E**), specifically with relation to PDAC_3 cells with multiple myeloid, CAF, and endothelial cell populations and their spatial localizations (**Supplementary Figure 15-17**). Notably, SEMA3C is upregulated in PDAC and correlates with poorer survival and increased invasiveness^39^.

### Stroma-derived CXCL12 & IL16 shape immune-excluded neighborhoods of persister populations

To benchmark SpaCCI and explore immunoregulatory LR circuits in human PDAC, we analyzed six spatial-transcriptomic specimens profiled with Xenium^40^: 3 treatment-naïve tumors and 3 tumors exposed to ChemoRT. High-resolution LR mapping revealed a prominent CXCL12– CXCR4 axis that is spatially confined to the stromal compartment (**Fig. 6A**). CAFs formed concentric rings of CXCL12 expression around malignant nests, coinciding with an abrupt loss of both CD8+ and CD4+ T cells at the tumor–stroma interface. Conversely, IL16 was enriched in myeloid clusters, and its gradient was accompanied by local accumulation of CXCR4-positive CD4+ T cells **(Fig. 6B**). Visium spatial visualization utilizing iSTAR further corroborated these patterns with malignant cells and T effector cells occupying separate discrete neighborhoods with greater overlap amongst iCAFs and ALT-TAMs (**Fig. 6C**). Quantitatively, CAF-derived CXCL12 and T-cell CXCR4 were both significantly up-regulated after ChemoRT (adjusted P < 0.0001; **Fig. 6D**), paralleled by an increased median T-cell–cancer-cell distance and a reduced T-cell–CAF distance (**Fig. 6E**). Neighborhood profiling centered on non-responder malignant states (PDAC_3) showed a marked rise in CXCL12 and IL16 compared with responder states (PDAC_0/1/2 & PDAC_prolif) (**Fig. 6F**). Stratifying the tissue into epilesional, juxtalesional and perilesional zones underscored this immune exclusion: cytotoxic CD4_c4_CTL cells and multiple CD8 subsets were depleted in epilesional areas but progressively re-appeared toward the periphery, with the effect accentuated in ChemoRT specimens (**Fig. 6G–I**). Collectively, these data indicate that RT intensifies stroma-driven CXCL12-mediated T-cell exclusion and IL16 recruitment of CXCR4+ CD4+ T cells, establishing an immune-suppressive niche that is most enriched in PDAC_3 neighborhoods.

**Figure 6:**
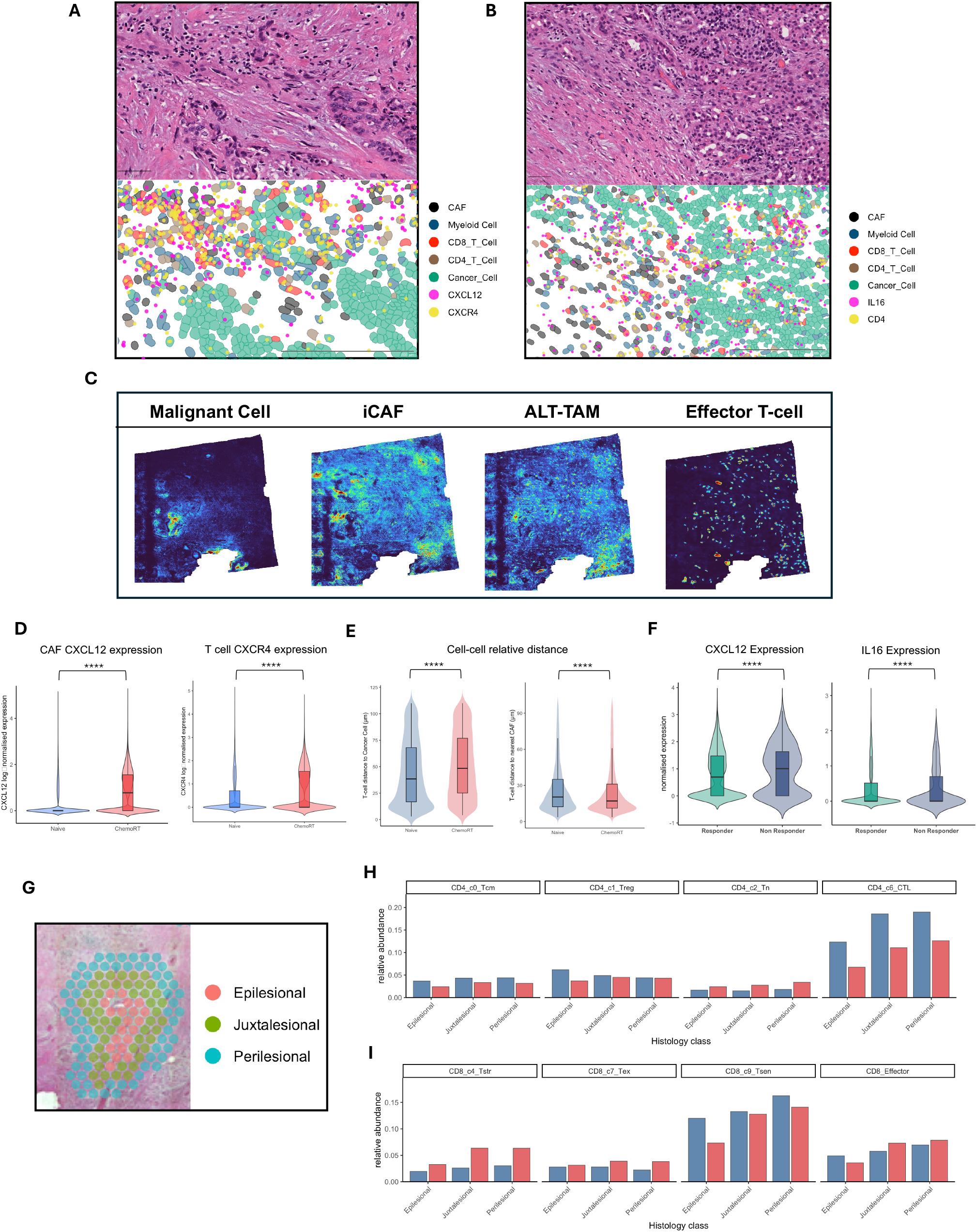
H&E of PDAC tissues with associated Xenium spatial map demonstrating colocalization of CXC12-CXCR4 (A) and IL16- CD4 signaling (B) ligand receptor interactions leading to immune exclusion zones. C Representative iSTAR signature scores of malignant and stromal populations demonstrating spatial localization of cells of interest. D. Bar plots of CXCL12 and CXCR4 normalized expression within CAF and T cell populations at treatment naïve status and following ChemoRT derived from Xenium (p<.0001). E. Bar plots of T cell to cancer cell (left) and CAF (right) distances following chemoRT derived from Xenium (p<.0001). F. CXCL12 and IL16 expression within surrounding stromal neighborhoods of responder and non-responder (persister cells) cancer cell subtypes derived from Visium (p<.0001). G. Representative visual of spatial spot selection methodology of epilesional (tumor bed), juxtalesional, and perilesional areas within tissues. H. Relative abundance of cell types of interest within spatial localizations based on Visium in treatment naïve (blue) and ChemoRT (red) tissues.

## Discussion

Our single-cell and spatial transcriptomic analyses revealed profound RT-induced remodeling in both malignant and non-malignant compartments of PDAC. Post-RT tumors showed selective enrichment or depletion of specific cell populations. For example, expansion of apCAFs and myCAFs suggested that RT favors fibroblast phenotypes associated with immune regulation and tissue repair. Similarly, we observed changes in immune cells, including an increase in ALT-TAMs and a redistribution of CD8+ T cell subclusters (notably the CD8_c4_Tstr subset). In the malignant compartment, most tumor cells subtypes were sensitive to RT; however, a distinct PDAC_3 cell state persisted. This surviving PDAC_3 subtype exhibited a unique metabolic expression program characteristic of defense against cuproptosis and ferroptosis. SpaCCI further demonstrated that spatial neighborhoods of these persister cells were enriched in CXCL12 and IL16 expression with associated immune cell exclusion.

The SpaCCI analysis framework integrates LR expression with spatial proximity to infer cell communication networks. SpaCCI outperforms existing methods by imposing realistic distance constraints on potential interacting cells. Whereas other approaches often consider cell pairs that are far apart or rely on arbitrary distance cut-offs SpaCCI restricts analysis to neighboring cells, thereby capturing true paracrine and juxtacrine signals with higher specificity. Using SpaCCI, we uncovered RT-driven changes in intercellular signaling that were not apparent by transcriptomics alone. Notably, RT shifted fibroblast signaling toward a dominant TGFβ- and PDGF-enriched program, consistent with a wound-healing, immune-exclusionary stroma. TGFβ becomes strongly upregulated in irradiated fibroblasts^41^, leading to fibroblast activation and secretion of factors that increase tissue fibrosis and suppress immune responses. In this respect, we found that CD8+ T cells in treated tumors skewed toward a TGFβ-influenced tissue-resident/stressed profile (CD8_c4_Tstr) rather than an effector phenotype. SpaCCI also revealed a surge of CAF-secreted CXCL12 after RT, with matched up-regulation of its receptor CXCR4 on T cells. This chemokine axis is known to repel effector lymphocytes from the tumor bed and to polarize macrophages toward tumor-supportive phenotypes as supported by our findings^42^. The expansion of an immunosuppressive TAM program after RT suggests that macrophages are being “educated” by the post-RT milieu to support tumor regrowth and dampen immune response. This is in line with reports that high-dose RT can drive influx of pro-tumor, M2-polarized TAMs that disable T cell mediated responses^43^. Notably in this context, TAMs are intrinsically radioresistant^44^ and can rapidly infiltrate irradiated sites. We also detected RT-induced enrichment of IL-16 in tumor-associated macrophages, forming local gradients that attracted and retained CD4+ T cells. Together, these changes illustrate a complex immune reprogramming in which it is important to note the context of timing where RT may transiently boost antigen release and T cell priming, but by the time of tumor resection (12 weeks post-RT in our study), the immune landscape appears tilted toward suppression rather than sustained anti-tumor activity. Semaphorin signaling also emerged as a significantly enriched pathway after RT. Semaphorin 3 ligands (SEMA3C) were elevated in treated tumor regions, along with their receptors (plexins/neuropilins) on neighboring cells. These axon guidance pathways are critical signaling mechanisms in PDAC ^37, 45^, influencing immune evasion, therapeutic resistance, perineural invasion, and stemness. However, their role in modulating RT response remains poorly defined. Further characterization of these pathways may uncover novel therapeutic vulnerabilities.

In the malignant epithelial compartment, our analyses identified a therapy-resistant tumor cell population with distinct metabolic adaptations. Our finding that the persisting tumor cells (PDAC_3) harbor metabolic dependencies linked to ferroptosis and cuproptosis resistance suggests that RT selects for cells with robust redox homeostasis and heavy-metal detoxification capacity. Gene signatures of ferroptosis resistance were prominent in PDAC_3, including elevated glutathione and lipid peroxide detoxification pathways. This result complements recent mechanistic studies showing that RT can induce ferroptotic cell death in cancer cells, which may function as a synergistic therapeutic node^46, 47^. The role of cuproptosis in RT-induced cell death has also been recently established^48^, with our data further supporting the role of metabolic plasticity (e.g. upregulation of thiol-rich metallothioneins or alterations in mitochondrial metabolism) may protect persister cells from copper-catalyzed toxicity. Notably, cuproptosis regulators have recently been implicated in PDAC progression and therapy response, hinting that targeting this vulnerability could be an unexplored opportunity ^49, 50^. Altogether, the persistence of the PDAC_3 subtype underscores a potential population of cells with specific resistant mechanisms that may be overcome with rational combinatorial RT regimens.

While this study provides a comprehensive view of PDAC treated with RT, we acknowledge several limitations. First, our single-cell RNA-seq dataset compared pre-RT vs. post-RT samples (largely capturing the effect of radiation alone), whereas our spatial transcriptomics compared untreated vs. post-*chemo*RT samples. The lack of a dedicated “chemo-only” control means that some changes observed in the spatial data could be partly due to chemotherapy. As a result, attributing all changes exclusively to RT must be done with caution. Second, the post-therapy samples were obtained approximately three months after completing RT. This relatively long interval may have allowed transient effects to subside and chronic adaptations to take hold. We might have missed short-lived waves of immune activation or acute tumor cell senescence that occur soon after irradiation. Longitudinal sampling at multiple time points would be needed to fully map the kinetic sequence of changes. Lastly, sample size and inherent variability in human tissues impose limitations on statistical power for some rarer cell populations and interactions. We attempted to mitigate sample to sample variability by focusing on findings that were consistent across both datasets, but certain trends that are unique to either one dataset cannot be excluded as being functionally relevant in the context of RT.

In summary, this study provides a comprehensive view of how PDAC tumors and their microenvironment adapt to radiation therapy. We document a shift toward fibroblast-driven fibrosis and immunosuppression, an immune landscape skewed by suppressive macrophages and stressed T cells, and the emergence of a metabolically reprogrammed tumor cell subset capable of withstanding therapy. These changes, validated through spatial profiling, deepen our understanding of PDAC biology under therapeutic stress and highlight tangible targets for intervention.

## Materials and Methods

### Patient sample collection

We conducted single-cell RNA sequencing (scRNA-seq) on 16 PDAC biopsies of matched pre- and post-treatment samples from a Phase I/II dose-escalation clinical trial of pancreatic stereotactic body radiation (SBRT). Additionally, Visium CytAssist spatial transcriptomics (ST) (10X Genomics) was performed on primary PDAC FFPE tissues collected at surgery from a separate cohort of patients (n=28; 8 untreated, 20 treated with chemoradiation). All biospecimens were collected under Institutional Review Board–approved protocols and obtained using written informed consent. At MD Anderson Cancer Center, tissue samples were collected under the Pancreas Tissue Bank protocol LAB00-396 and subsequently analyzed under a use protocol PA15-0014. An additional 6 PDAC human tissue specimens (3 untreated, 3 treated with chemoradiation) profiled with Xenium spatial transcriptomics using a custom gene panel were obtained from an external dataset^40^. Tissues selected for spatial transcriptomics were reviewed by a gastrointestinal pathologist and included for analysis if they contained at least 20% viable tumor tissue. This study was conducted in accordance with Good Clinical Practices concerning medical research in humans per the Declaration of Helsinki.

### Single cell RNA-sequencing (scRNA-seq) and data analysis

scRNAseq libraries were generated by 10X Chromium System (10X Genomics). Samples were run on the Illumina NextSeq500 (Illumina). For data analysis, we used Cell Ranger, version 3.0.1 (10X Genomics) to process raw sequencing data. After sequencing and data processing, downstream analyses were performed using Seurat (v4.4.0) package in R (v4.3.1). Single cells were clustered with FindClusters function and manually annotated to known cell types using known established cell markers. FindAllMarkers with Wilcoxon Rank Sum test in Seurat was utilized to identify the markers genes for each cell type(min.pct>0.25 and logfc.threshold>0.25). Enriched pathways for lists of marker genes were determined by employing the gene set collections at the Molecular Signature Database (Broad Institute, USA) using a hyper-geometric distribution (P < 0.05). Raw counts from each sample and cell type were summed up to generate pseudocounts, which was analyzed with DESeq2^51^ to identify significantly differentiated genes between groups in each cell type. Gene sets enriched in each group is identified utilizing the GSEA method^52^, based on 1000 random permutations of the ranked genes. Changes in cell type abundances between groups were assessed using a binomial generalized linear model (GLM) with emmeans package.

### Spatial transcriptomics and data analysis

After sectioning of FFPE blocks onto the Visium slides, sample processing was performed as described in the Visium CytAssist 10x Genomics. After library preparation, samples were pooled and sequenced using a NextSeq 550 (Ilumina).The sequencing outputs for all Visium datasets were preprocessed using SpaceRanger (v2.1.0) and mapped to the GRCh38 human reference genome, and the filtered matrices were converted to Seurat objects in R and used for downstream analyses. Cell type composition in ST samples were inferred with Cytospace, RCTD, and iSTAR^12- 14^ using the annotated single cell object.

### TCGA-PDAC deconvolution and survival analysis

RSEM counts for transcriptomics data from TCGA Pancreatic Adenocarcinoma patients treated with radiation were downloaded from Broad Institute^53^. TCGA counts were deconvoluted to cell types found in the single cell data using BayesPrism (v2.2.2)^54^. Survival analysis was performed on the radiation treated patients after separating the samples according to cell composition of the desired cell type.

### Pancreatic orthotopic injection tumor model

0.25 million of AMKPC-Luc cell (isolated from Trp53^R172H/+^, LSL-Kras^G12D^, Pdx-1-Cre mouse with C57Bl/6J background) was suspended in 20uL DPBS and orthotopically injected to C57Bl/6J mice pancreas by BD 29G insulin syringe under the sterile surgical condition. One titanium microclip (WECK Horizon 005200) was clipped 1mm under the injection point to visualize the tumor site under the integrated CT on Small Animal Radiation Therapy Platform (SARRP). 10 days were given to allow the wound recovery and tumor establishment before the tumor volume was measured by IVIS and the mice were divided into treatment groups.

### Mouse Stereotactic Body Radiation Therapy (SBRT)

SBRT treatment group mouse is anaesthetized with 3% isoflurane and restrained on the treatment platform in SARRP. 360-degree CT scan is performed to visualize the titanium microclip. The radiation isocenter was set on 5mm above the microclip vertically and 8 Gy X-ray was delivered to the isocenter from four angles (5°, 30°, −175°, and −150°), repeated daily for five consecutive days.

### Collection of recurrent tumors and establishment of relapsed tumor cell lines

At the primary endpoint (animal is moribund or tumor size reaches to IACUC standard), the tumor bearing mice were euthanized and the tumors were isolated. Tissue was recovered and digested in Hank’s balanced salt solution with 1x DNase I and Collagenase IV for 1 hours. The cell suspension was then filtered via a 70-um cell strainer and resulting single cell suspension proceeding to single cell RNA sequencing utilizing the 10X Chromium Single Cell 3’ GEM v3.1 protocol as described above or seeded into 10-cm cell culture dish with completed DMEM (10% (volume/volume; v/v) heat-inactivated FBS (GE HyClone) and 1% (v/v) penicillin/streptomycin (Gibco) at 37 °C and 5% CO2.

### Cancer cell lines

All cells were cultured in a 37 °C incubator in an atmosphere of 5% CO_2_. All cells were cultured in Dulbecco’s modified Eagle medium supplemented with 10% fetal bovine serum and 10,000 U/mL of penicillin-streptomycin.

### SpaCCI method overview

SpaCCI is a cell-cell interaction detection tool that primarily design for spot-level spatial transcriptomics, where it takes both cell type mixture and cell type abundance into account. In general, SpaCCI requires two types of data inputs: (1) spot-level spatial transcriptomics data, which includes the gene expression profiles and spatial coordinates, and (2) cell type proportion data, which can be obtained through existing methods such as Robust Cell Type Deconvolution (RCTD) or conditional autoregressive-based deconvolution (CARD). Additionally, a selected ligand-receptor database is used as a reference. The primary goal of SpaCCI is to detect significant spatial cell-cell interactions that may occur within the spatial domain at local, regional, and global levels. The SpaCCI software package is publicly available at https://litingku.github.io/SpaCCI/.

#### Non-negative least square regression

Let *β*_*g*_= (*β*_{1,*g*}_, …, *β*_{_*K*,*g*_}_)^*T*^ be the non-negative vector of the average gene expression of a gene *g* in *K* cell types in the region *S* of a tissue slide, with *n* spots in this region. We use the non-negative least square regression to estimate how different cell types contribute to the gene expression in this region using cell type proportion matrix as the predictor matrix.

Consider a linear regression model:

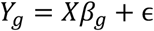

where *Y*_*g*_= (*y*_{1,*g*}_, …, *y*_{*n*,*g*}_)^*T*^ denote the observed gene expression of a gene *g* across all *n* spots in region *S* and 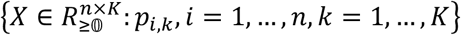 a *n* by *K* matrix denote the cell type proportion matrix of all *K* cell types across all *n* spots and *p*_*i*,*k*_ represent the proportion of cell type *k* in spot *i*.

We estimate 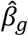 by

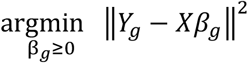

then estimated gene expression of a gene *g* for all cell types *K* in the region *S* is 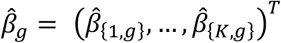 denote as the gene expression of a gene *g* for all cell types *K* in the region *S*.

### *Raw ligand-receptor interaction* strength

Consider a ligand-receptor pair, where the ligand *L* consists of *l* genes (*i. e*., *L*_1_, . ., *L*_*l*_), the receptor *R* consists of *r* genes (*i. e*., *R*_1_, . ., *R*_*r*_).

The raw gene expression of ligand *L* and receptor *R* in all cell types *K* are estimated by first, using non-negative least square regression to obtain the gene expression of each gene and second, approximates by the geometric mean if the ligand or the receptor consists of multiple genes. Finally, inspired by CellChat, we take the cell type proportion into consideration and use a Hill function to model the interaction strength between a ligand-receptor pair with a parameter *K*_*h*_ whose default value was set to be 0.5. The interaction strength P_*i*,*j*_ between cell type *i* and cell type *j* of a ligand-receptor pair *N* is calculated as

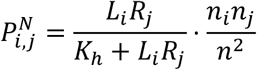

where *L*_*i*_ denoted as the raw gene expression of a ligand *L* in cell type *i, R*_*j*_ denoted as the raw gene expression of a receptor *R* in cell type 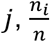 denoted as the mean cell type proportion of cell type 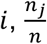 denoted as the mean cell type proportion of cell type *j*.

### Permutation test for regional detection

We use the permutation test to access whether the interaction strength of the ligand and the receptor expressed significantly in a given region *S* for *K* cell types compare to the rest of the regions on the tissue slice, denoted by 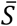. We define the null hypothesis and alternative hypothesis as

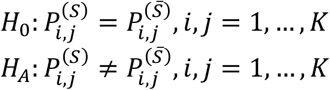

where 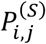 represents the interaction strength of the ligand *L* in cell type *i* and the receptor *R* in cell type *j* in the region *S*, and 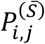 represent the interaction strength of the ligand *L* in cell type *i* and the receptor *R* in cell type *j* outside the region *S*.

To perform pairwise comparisons between all cell types simultaneously, we employed a random permutation strategy, shuffling both the cell type labels across the dataset. This approach was repeated to recalibrate the expressions of ligands and receptors via our non-negative least squares regression model. Specifically, our permutation protocol involves considering not only the spatial locations of the spots but also the cell type labels within the cell type proportion matrix. Crucially, we ensure that each cell type label is reassigned to a proportion from a different spot, preventing any cell type from retaining its original cell type label proportion. This rigorous shuffling method helps to generate a robust null distribution under the null hypothesis, essential for our statistical testing framework.

For the *p*-th permutation, we compared the permuted interaction strength for each ligand-receptor pair as 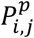 to the raw (null) interaction strength P_*i*,*j*_ calculated initially for all cell types *K* in a region *S*. If the permuted interaction strength was greater than the raw (null) interaction strength, we denoted this with 1, indicating a more extreme value under the permutation than observed, and 0 otherwise.

With the binary indicator, the p-values were calculated for cell types *i, j* = 1, …, *K* by

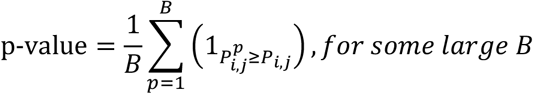

To control the false discovery rate (FDR) for pairwise comparisons across all cell types, we apply Benjamin-Hochberg (BH) method, which controls the expected proportion of falsely rejected null hypotheses among all rejected hypotheses. This allows us to identify cell types with significantly different ligand and receptor expressions while minimizing the risk of false discoveries due to pairwise comparisons.

### Permutation test for localized detection

To identify localized hot spots of cell-cell interaction across the entire slide, we employed a permutation framework similar to our regional approach but tailored to detect localized specificity. Each spot’s small neighborhood was defined with a 200 µm interaction range, considered the critical distance for significant biological interactions.

For robust assessment of cell-cell interactions within these defined neighborhoods, we implemented a random permutation strategy. This involved systematically shuffling cell type labels across various neighborhoods. Such recalibration of ligand and receptor expressions was facilitated using our non-negative least squares regression model, and the interaction strength is calculated considering cell type abundance and Hill function. This localized detection strategy was applied consistently to all small neighborhoods across the slide, allowing for comprehensive permutation testing. The methodology ensures robust inference of interaction hot spots throughout the slide, capturing both the microscale intricacies and the broader spatial patterns of cellular communication.

### Permutation test for global detection

To assess cell-cell interactions from a global perspective, we utilized a permutation framework similar to that used for regional detection. We define the null hypothesis *H*_0_ as the interaction strength between any given ligand-receptor pair is consistent across all cell types on the whole slide, whereas the alternative hypothesis as the interaction strength between ligand-receptor pair varies among the cell types across the whole slide.

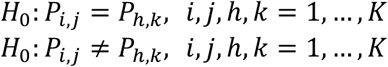

where P_*i*,*j*_ represents the interaction strength of the ligand *L* in cell type *i* and the receptor *R* in cell type *j* in the whole slide, and P_*h*,*k*_ represent the interaction strength of the ligand *L* in cell type *h* and the receptor *R* in cell type *k* outside the whole slide.

This global analysis allows us to understand broad patterns of intercellular communication, offering insights into the homogeneity or heterogeneity of cellular interactions across the whole slide and different cell types.

### Simulation Setting using Mouse Brain Stereo-seq Data

#### Mouse Brain Stereo-seq Data

We downloaded the adult mouse brain Stereo-seq data from Chen A, Liao S, Cheng M, et al. [https://db.cngb.org/stomics/mosta/]. In total, there are 34,263 effective bins with a mean of 3,561 UMIs and 1,335 genes detected per bin.

#### Simulated datasets

To ensure that the cell-cell interaction analyses of the Stereo-seq dataset were comparable across SpaCCI, CellChat, and CellPhoneDB, we simulated the spot-level spatial transcriptomics (ST) dataset for SpaCCI using the adult mouse brain Stereo-seq data. Initially, we used the spatial coordinates from the Stereo-seq dataset to group neighboring single cells into spots by setting a bin size of 5, resulting in a total of 1,625 spots across the adult mouse brain, while preserving the original spatial coordinates. We then computed the spot-level gene expression profiles by aggregating the expression data from the selected single cells within each spot. Regarding the spot-level cell type proportions, the original Stereo-seq dataset contained 20 distinct cell types. To simplify the analysis and enhance interpretability, we performed hierarchical clustering to reduce the number of cell types, ultimately grouping them into 8 major cell types. This approach allowed us to create a simulated spot-level ST dataset that closely mirrors the original adult mouse brain Stereo-seq data, facilitating a robust comparison of cell-cell interaction detection capabilities among SpaCCI, CellChat, and CellPhoneDB.

#### Compared methods for cell-cell interaction detection

As Stereo-seq technology combines single-cell resolution with spatial transcriptomics, we input the original adult mouse brain Stereo-seq data into CellChat and CellPhoneDB, both of which tools are primarily designed for single-cell resolution ST datasets. To ensure a fair comparison, we used the simulated spot-level dataset from the Stereo-seq data and analyzed it using SpaCCI. We then compared the results across these methods, evaluating the global cell-cell interaction patterns by calculating the Pearson correlation between the results, thereby assessing the consistency and accuracy of SpaCCI relative to CellChat and CellPhoneDB.

#### Localized detection accuracy

To assess the accuracy of SpaCCI in detecting localized cell-cell interactions, we first calculated the interaction strength across the entire tissue slide using the original Stereo-seq dataset, following the method outlined in the “Raw Ligand-Receptor Interaction Strength” section. This served as the ground truth for the simulated spot-level data. Using SpaCCI, we then identified localized cell-cell interaction hotspots by outputting the interaction probability for each spot. With both the ground truth and the calculated probabilities, we plotted the average ROC curve for the ligand-receptor interactions across various combinations of cell type pair interaction directions.

### Simulation Setting using Human Pancreatic Single Cell Data

#### Human Pancreatic Single Cell Data

We downloaded the count matrix along with cell type annotations from Baron M, Veres A, Wolock SL, et al. [https://www.ncbi.nlm.nih.gov/geo/query/acc.cgi?acc=GSE84133](GSE84133). The dataset includes UMI counts and annotations for 14 cell types, including alpha, beta, delta, gamma, epsilon, acinar, ductal, activated stellate, quiescent stellate, endothelial, macrophage, mast, T cells, and Schwann cells. On average, 1,049 UMIs and 20,215 genes were detected per cell.

#### Simulated datasets

To evaluate the performance of SpaCCI in detecting cell–cell interactions, we generated a simulated spot-level spatial transcriptomics (ST) dataset within an in silico tissue environment. This simulation accounted for cell type-specific gene expression distributions and cell type proportions, allowing us to test assumptions regarding the impact of cell type heterogeneity and the occurrence of false positives in cell–cell interaction analyses. We selected six cell types—beta, delta, ductal, activated stellate, quiescent stellate, and macrophage—based on their distinct expression profiles in the original scRNA-seq dataset. The simulation process began by estimating the gene expression distribution for each gene within these cell types using a negative binomial distribution. This in silico tissue was designed to contain 782 spots, each comprising 12 single cells, and was organized into six regions. For each cell type, gene expression was estimated based on its abundance in the reference scRNA-seq dataset, focusing on 13 selected ligand/receptor genes. To maintain the sparsity characteristic of scRNA-seq data, the proportion of zero counts for each gene was calculated and incorporated into the simulation. Additionally, we introduced spatial heterogeneity by randomly assigning higher expression levels to the central region and lower expression levels to the peripheral regions.

#### Compared methods for cell-cell interaction detection

We compared the performance of SpaCCI with two widely used existing methods, CellChat and CellPhoneDB, ensuring that the ligand-receptor interactions present in our simulated datasets were included in both databases. For regional cell-cell interaction detection, we generated 100 simulated datasets, each containing a ground truth consisting of 27 ligand-receptor pair interactions occurring specifically in the central region. These datasets were then analyzed using SpaCCI, CellChat, and CellPhoneDB. To evaluate the performance of each method, we employed a binary indicator approach, where a value of 1 indicated that the interaction was successfully detected, and a value of 0 indicated no detection. From these binary results, we calculated the average sensitivity (the proportion of true positive interactions correctly identified) and specificity (the proportion of true negative interactions correctly identified) across all simulated datasets. For global cell-cell interaction detection, we focused on the incorrect detection rate. Specifically, we calculated the probability of false positive detections in each simulated dataset. This measure allowed us to assess the robustness of each method in avoiding spurious detections across the entire spatial domain.

### Real data analyses

We applied SpaCCI to analyze published IPMN 10X Visium ST datasets from Sans M, Makino Y, Min J, et al. [https://www.ncbi.nlm.nih.gov/geo/query/acc.cgi] (GSE233254) and the PDAC 10X Visium ST datasets studied in this paper. Both datasets provide cell type proportions derived from RCTD, and we utilized the CellChat ligand-receptor database as the reference for our analyses.

### Regional cell-cell interaction detection

For the IPMN 10X Visium ST datasets, the study annotated three distinct regions based on research interests: juxtalesional, epilesional, and perilesional regions. We focused on the juxtalesional region to compare the regional cell-cell interaction patterns among the low-grade (LG) IPMN sample, high-grade (HG) IPMN sample, and IPMN-associated cancer sample (PDAC). This comparison aimed to uncover interaction pattern differences associated with IPMN progression. The analysis was conducted following the SpaCCI regional detection protocol. To validate our findings, we employed immunofluorescence (COMET) data, providing an additional layer of confirmation for the detected interactions.

### Global cell-cell interaction detection

For the PDAC 10X Visium ST datasets analyzed in this study, we compared the global cell-cell interaction patterns between the Naïve treatment and chemoradiation (chemoRT) treatment groups. Following the SpaCCI global detection protocol, we applied SpaCCI to each sample within both treatment groups. To facilitate comparisons, we first aggregated the interaction results for each treatment group. We then compared the interacting cell type pairs between the Naïve and chemoRT treatment groups using the Wilcoxon test. To examine differences in specific ligand-receptor interactions between the two treatment groups, we calculated the log2FoldChange of the sum of ligand-receptor interaction enrichment within each group. Statistical significance was assessed using a student’s t-test, with p-values adjusted for multiple comparisons using the Benjamini-Hochberg (BH) method. To visualize the results, we generated a volcano plot. Interactions were declared as upregulated in the chemoRT group if the BH-adjusted p-value was less than 0.1 and the log2FoldChange was greater than 0. Conversely, interactions were considered downregulated in the Naïve group if the BH-adjusted p-value was less than 0.1 and the log2FoldChange was less than 0.

### Local cell-cell interaction detection

For the PDAC 10X Visium ST datasets, we applied SpaCCI to detect localized cell-cell interaction hotspots within samples from each treatment group. To evaluate the spatial distribution and intensity of these interactions, we visualized the results by plotting the detected hotspots directly onto the spatial tissue slides. This approach allowed us to observe and compare the localized interaction patterns between the treatment groups, providing insights into how treatment may influence the spatial dynamics of cell-cell interactions within the tumor microenvironment.

## Acknowledgements

We gratefully acknowledge the late Cullen M. Taniguchi, MD, PhD, for his mentorship, enduring support, and invaluable contributions to this work; his example continues to inspire our efforts.

V.B was supported by the American Society of Clinical Oncology Young Investigator Award. A.M. was supported by the MD Anderson Pancreatic Cancer Moon Shot Program; the Sheikh Khalifa Bin Zayed Al-Nahyan Foundation; Break Through Cancer; and NIH grants (U54CA274371, U01CA200468 and U24CA274274). E.J.K. was supported by U54CA210181, U54CA143837, U01CA196403, U01CA200468, U01CA200468, U01CA214263, P50CA221707, R01CA221971, R01CA218004, P30CA016672, RP220119. L.K, L.L, and Z.L. were partially supported by U24CA274212. J.M. was supported by a Pilot/Feasibility Award from the Texas Medical Center Digestive Disease Center (P30DK056338) and a Seed Grant from the Hirshberg Foundation for Pancreatic Cancer Research.

## Code Availability

The SpaCCI software package and source code have been deposited at https://litingku.github.io/SpaCCI/. All scripts used to reproduce all the analysis are also available at the same website

