## Supplementary Figure for "A spatial atlas of chemoradiation therapy in pancreatic cancer identifies cellular and microenvironmental determinants of persister populations"

### Supplementary Figure 1

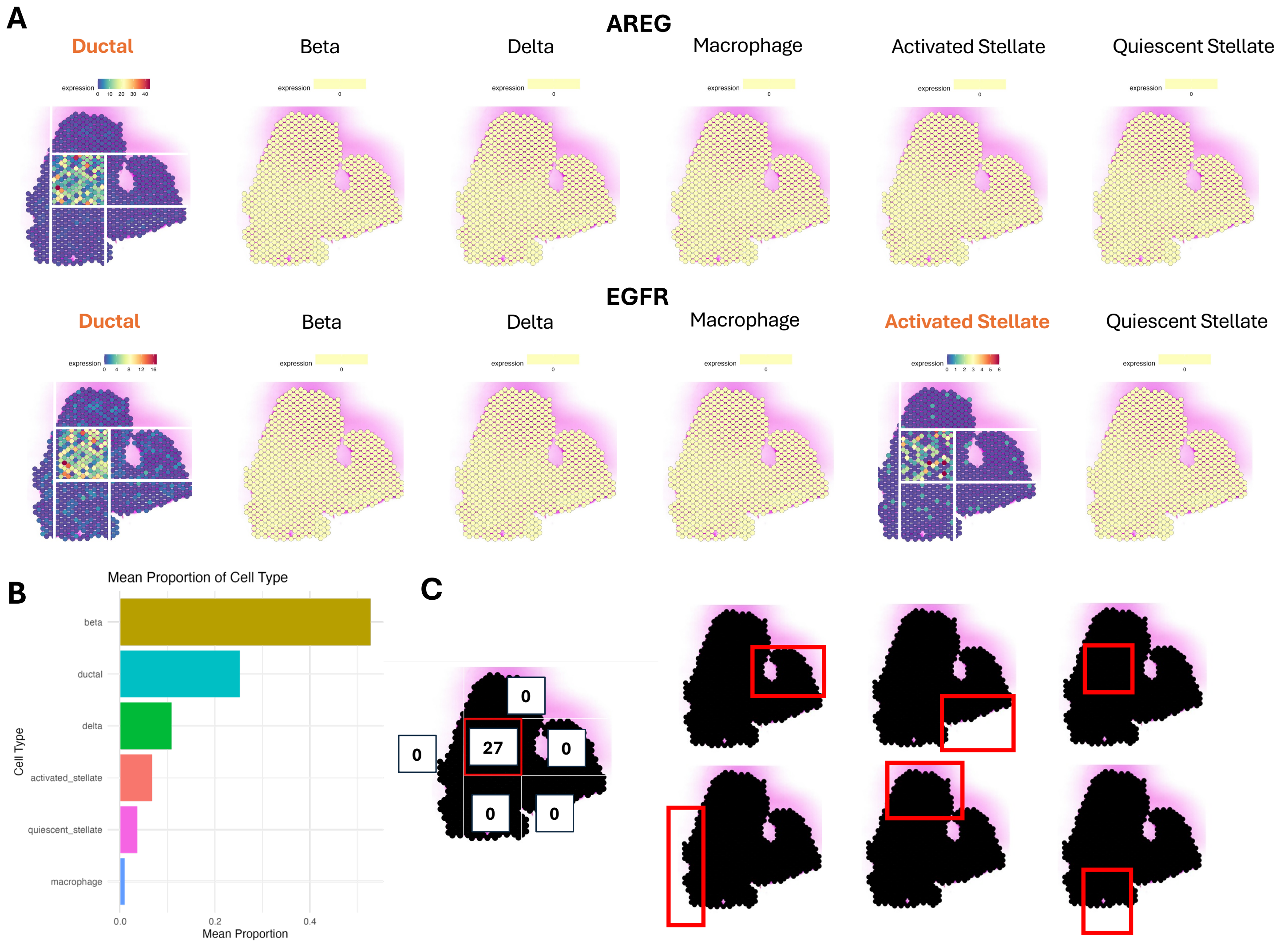


**Supplementary Figure 1**. Simulation design and evaluation framework for benchmarking regional cell–cell interaction detection. (A) Visualization of ligand (AREG) and receptor (EGFR) expression across six simulated cell types: Beta, Ductal, Delta, Macrophage, Activated Stellate, and Quiescent Stellate. The spatial distribution reflects ground truth localization, where interactions (e.g., Ductal → Ductal, Ductal → Activated Stellate) are engineered to occur in the central region of the tissue. (B) Bar plot showing the mean proportion of each cell type across simulated spatial slides. Beta cells dominate the tissue, while macrophages are rare, comprising less than 5% of the cell mixture. (C) Schematic of spatial region partitioning for evaluation. The whole tissue was divided into six regions, and 27 ligand–receptor interactions were specifically embedded in the center region to assess regional detection accuracy. This setup forms the basis for testing the ability of SpaCCI and other tools to localize interactions.

#### Supplementary Figure 2

**A**

**Expected CCI LR pair in the CENTER REGION compared to others**

| | Ligand | Receptor |
|---|---|---|
| CSF1 – CSF1R | ductal | macrophage |
| EDN1 - EDNRB | ductal | activated stellate |
| | ductal | quiescent stellate |
| EDN2 – EDNRB | ductal | activated stellate |
| | ductal | quiescent stellate |
| EDN3 - EDNRB | beta | activated stellate |
| | delta | activated stellate |
| | beta | quiescent stellate |
| | delta | quiescent stellate |
| EDN1 - EDNRA | ductal | activated stellate |
| | ductal | quiescent stellate |
| EDN2 – EDNRA | ductal | activated stellate |
| | ductal | quiescent stellate |
| EDN3 - EDNRA | beta | activated stellate |
| | delta | activated stellate |
| | beta | quiescent stellate |
| | delta | quiescent stellate |

| | Ligand | Receptor |
|---|---|---|
| AREG - EGFR | ductal | ductal |
| | ductal | activated stellate |
| TGFA - EGFR | ductal | ductal |
| | ductal | activated stellate |
| HBEGF - EGFR | ductal | ductal |
| | ductal | activated stellate |
| IGFBP3 – TMEM219 | ductal | ductal |
| | ductal | beta |
| | activated stellate | ductal |
| | activated stellate | beta |

**B**

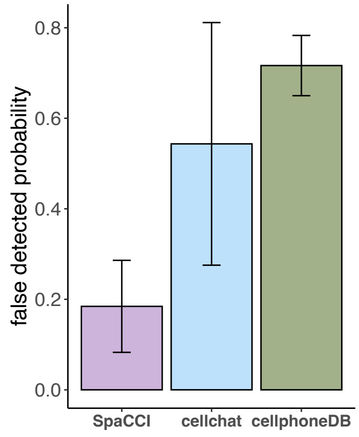


**Supplementary Figure 2**. Benchmarking dataset for assessing regional and global detection accuracy of cell-cell interactions. (A) Table of simulated ground truth ligand–receptor pairs, each defined with specific sender and receiver cell types. These interactions were embedded to occur specifically in the center region and used to evaluate directional detection accuracy. (B) Comparison of false positive detection probabilities across SpaCCI, CellChat, and CellPhoneDB over 100 simulation replicates. SpaCCI achieves the lowest false positive rate, highlighting its precision in global detection under cell type mixture conditions.

### Supplementary Figure 3

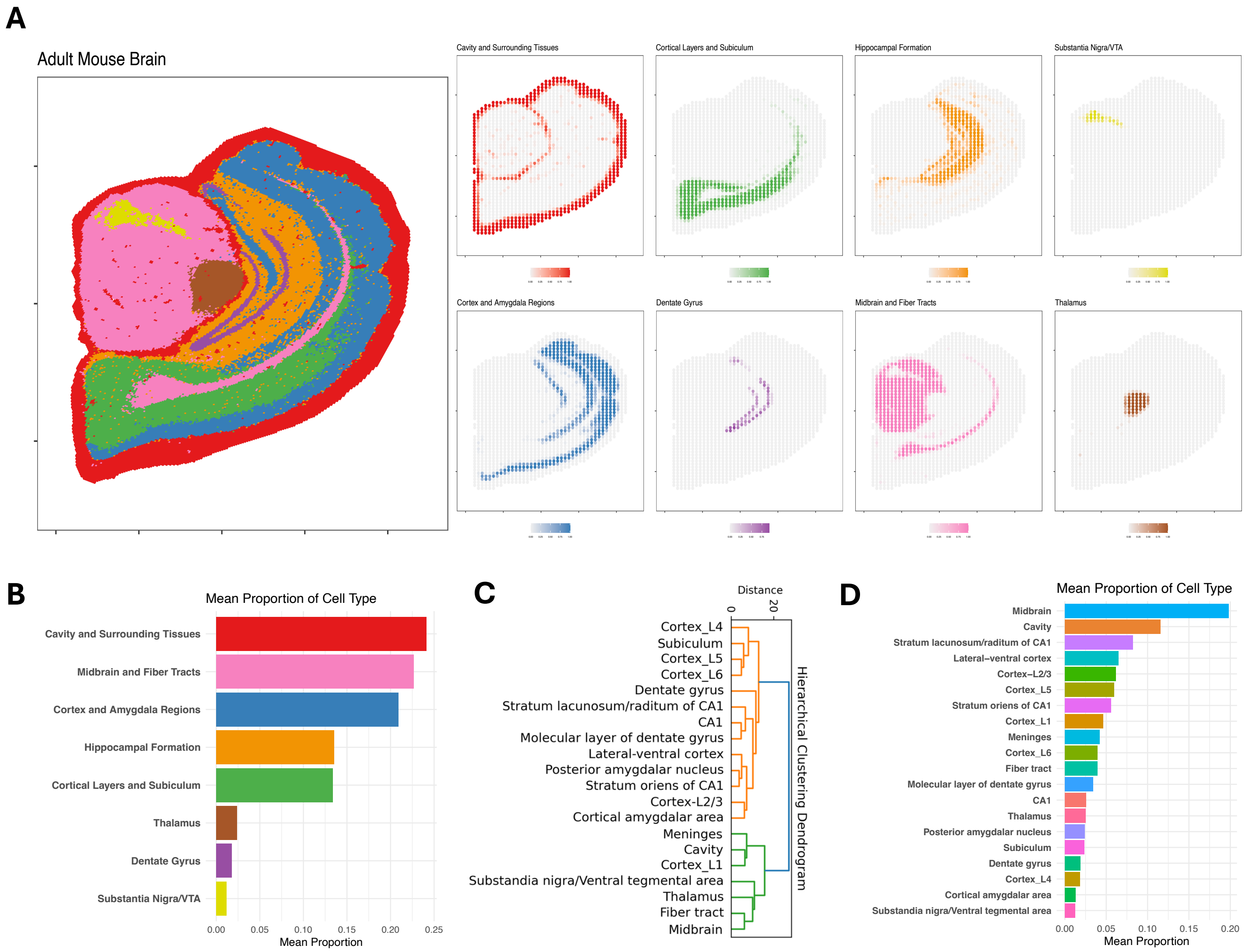


**Supplementary Figure 3**. Simulation of spot-level spatial transcriptomics data using adult mouse brain Stereo-seq. (A) Spatial annotation of the simulated brain tissue with eight major brain regions, derived from the original Stereo-seq dataset and aggregated for SpaCCI simulation. Cell type annotations and proportions were integrated with spatial coordinates to mimic the resolution and format of 10x Visium. (B) Bar plot showing the mean proportion of each major brain region across all simulated spots. The cavity and surrounding tissues and midbrain and fiber tracts made up the highest proportion, each exceeding 20%, while thalamus, dentate gyrus, and substantia nigra/VTA accounted for the lowest proportions (<5%). (C) Hierarchical clustering dendrogram used to group detailed original Stereo-seq cell annotations into the eight broader cell type categories used in the simulation. (D) Bar plot of the mean proportion of the original Stereo-seq cell types prior to hierarchical grouping, illustrating the diversity and abundance of brain cell populations.

### Supplementary Figure 4

**A**

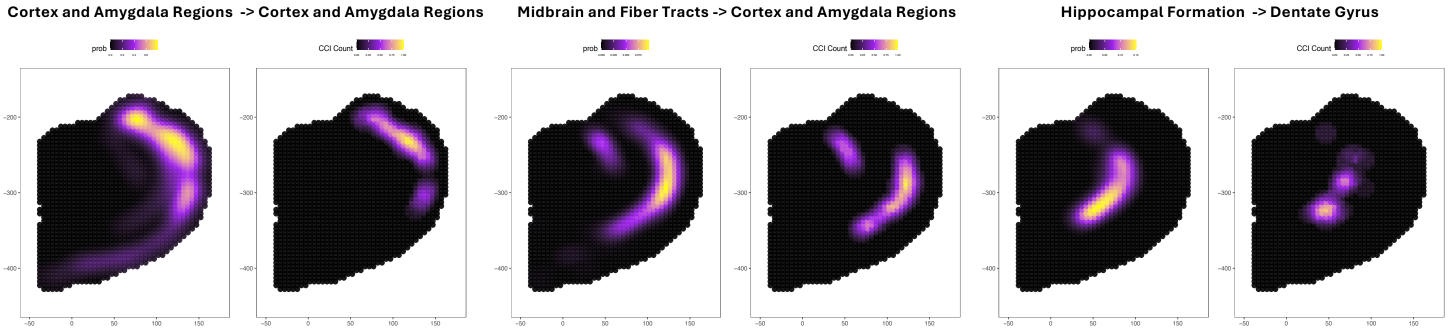


**Supplementary Figure 4**. Assessment of localized cell–cell interaction (CCI) detection accuracy using Stereo-seq mouse brain data. (A) Comparison of interaction hotspots between ground truth probabilities (left column in each pair) and SpaCCI-inferred CCI counts (right column in each pair) for three representative cell type pair combinations.

### Supplementary Figure 5

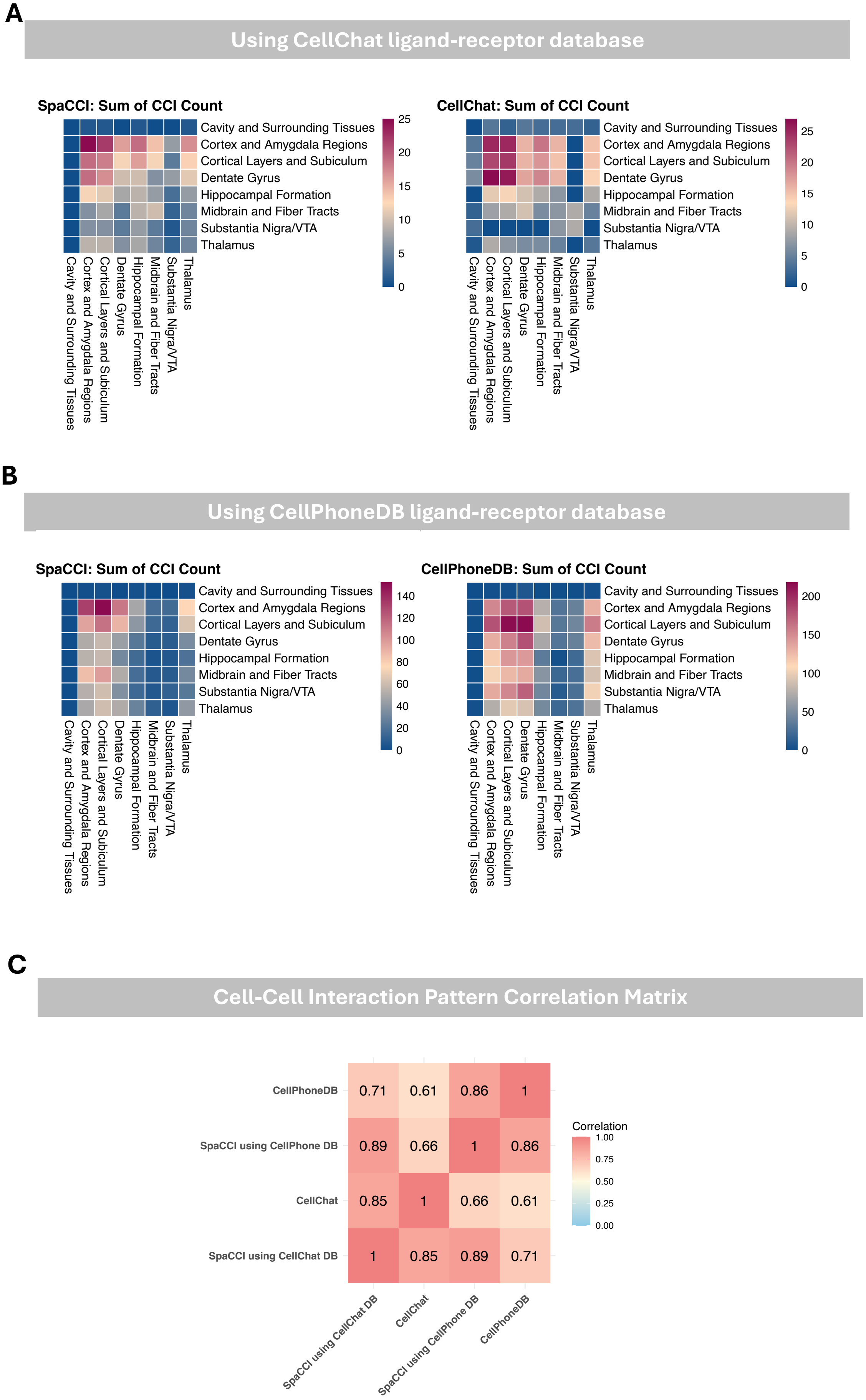

**Supplementary Figure 5**. Evaluation of global cell-cell interaction (CCI) pattern detection on Stereo-seq mouse brain data. (A) Heatmaps showing the sum of detected CCIs between brain anatomical regions using SpaCCI and CellChat, both based on the CellChat ligand–receptor database. Rows and columns correspond to anatomical regions annotated in the Stereo-seq dataset. (B) Similar analysis using the CellPhoneDB ligand–receptor database, comparing results from SpaCCI and CellPhoneDB. SpaCCI detects global CCI patterns consistent with those identified by CellPhoneDB, despite being designed for spot-level spatial transcriptomics. (C) Correlation matrix comparing the global interaction patterns inferred by SpaCCI (using both databases), CellChat, and CellPhoneDB. High correlation values (up to 0.89) demonstrate that SpaCCI produces consistent and robust global CCI patterns, comparable to single-cell-based methods, when applied to high-resolution Stereo-seq spatial data.

**Supplementary Figure 6**

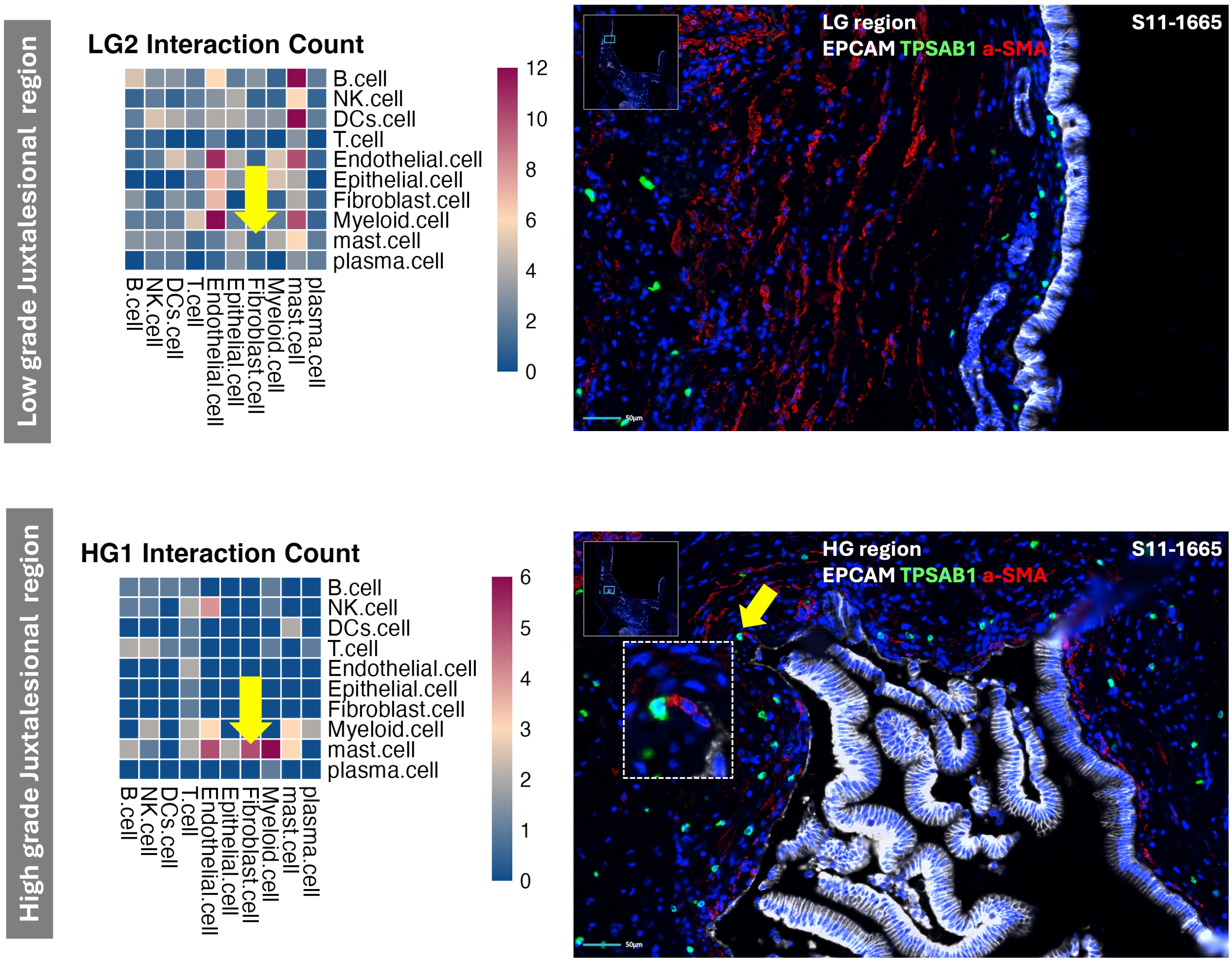


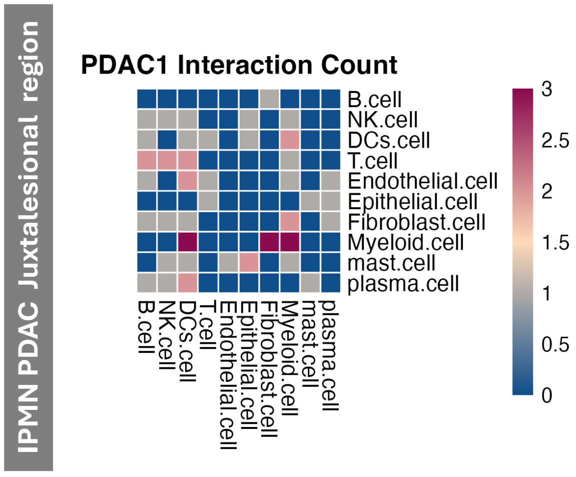


**Supplementary Figure 6**. Application of SpaCCI to 10X Visium spatial transcriptomics (ST) data from IPMN samples, with experimental validation using COMET immunofluorescence imaging. Three representative samples are shown: one low-grade (LG) IPMN (LG2), one high-grade (HG) IPMN (HG1), and one IPMN-associated pancreatic cancer sample (PDAC1). Left panels: Heatmaps display the number of significant cell–cell interactions identified by SpaCCI, with rows representing ligand-expressing (sender) cell types and columns representing receptor-expressing (receiver) cell types. Notably, mast cell–fibroblast interactions are significantly more frequent in the HG IPMN sample (HG1) than in the LG or PDAC samples. Right panels: COMET immunofluorescence staining validates this spatial pattern. In HG IPMN, mast cells and fibroblasts are frequently co-localized, supporting the SpaCCI-inferred interactions. In contrast, in the LG IPMN sample, mast cells are more spatially separated from fibroblasts.

#### Supplementary Figure 7

A

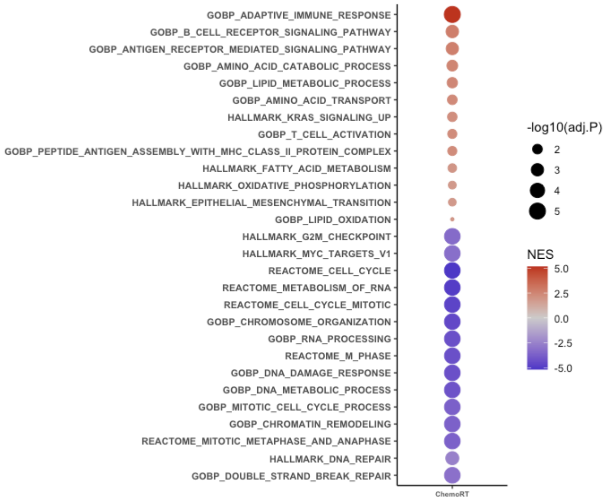


B

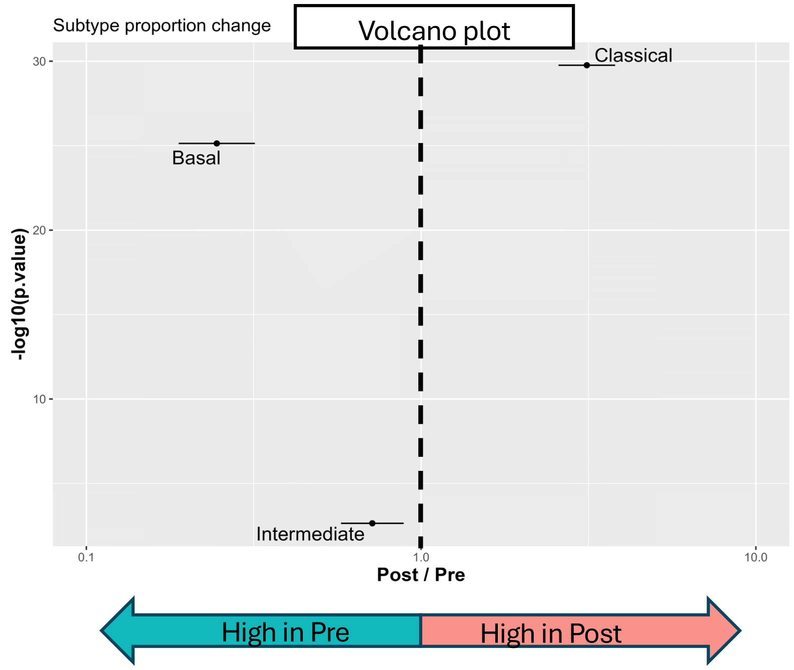


| Cell_Type | p.value | -log10(Pvalue) | odds.ratio | log2_odds.ratio |
|---|---|---|---|---|
| Classical | 1.73E-30 | 29.76 | 3.126080117 | 1.644 |
| Intermediate | 0.002293 | 2.64 | 0.714815435 | -0.484 |
| Basal | 7.39E-26 | 25.13 | 0.245260015 | -2.028 |

C

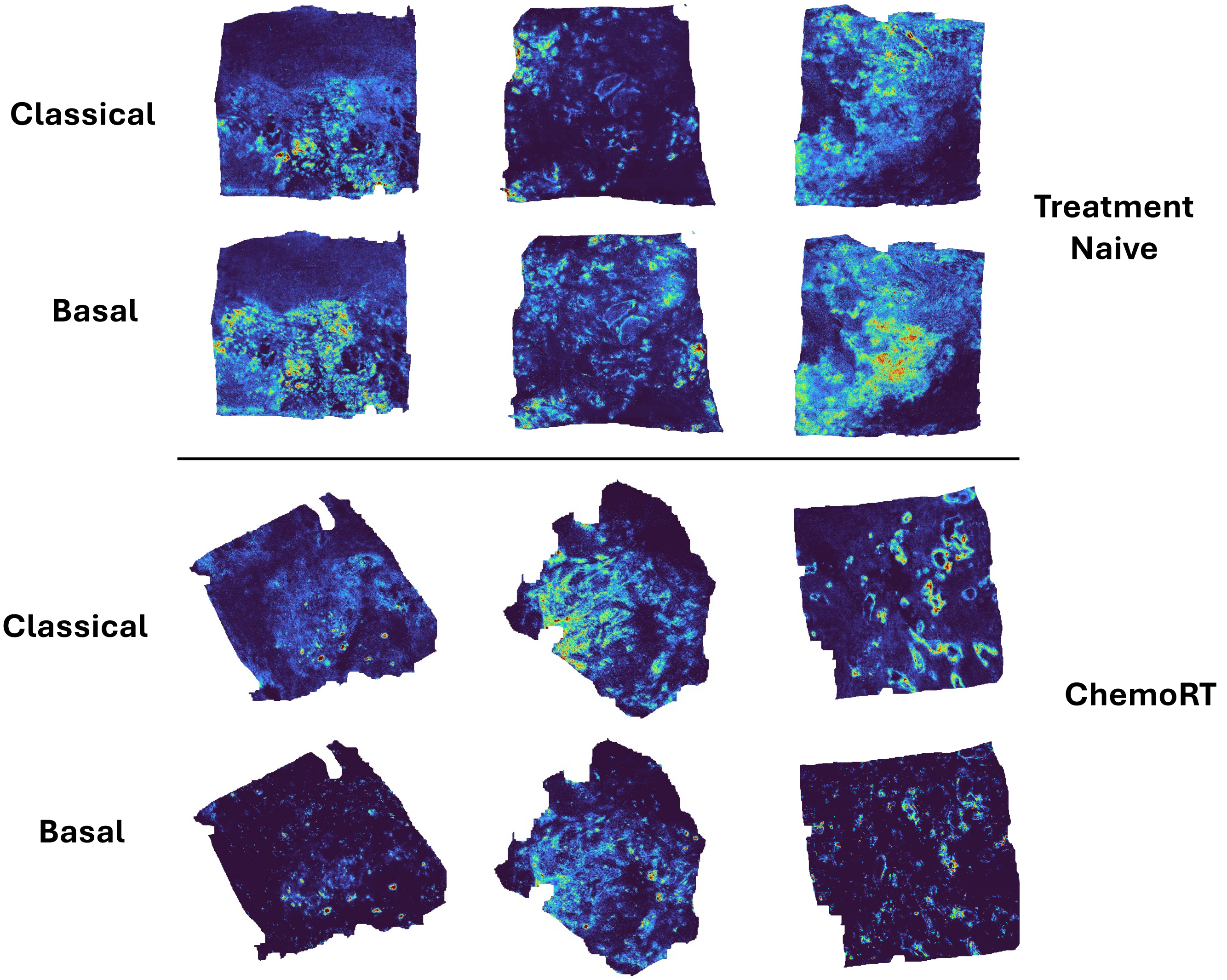

**Supplementary Figure 7**. A. Dotplot of significantly enriched upregulated (red) and downregulated (blue) pathways comparing pseudobulk expression of chemoRT vs Naïve tissues. B. Volcano plot displays odds ratios (Post / Pre) for Classical, Basal, and Intermediate PDAC subtypes based on single-cell RNA-seq of matched pre- and post-RT samples. Subtypes to the right of the dashed line are enriched post-RT; those to the left are enriched pre-RT. Summary statistics are shown in the accompanying table. C. Spatial visualization of classical and basal molecular subtypes imputed by inferring Super-resolution Tissue ARchitecture (iSTAR) from representative naïve and chemoRT treated samples.

**Supplementary Figure 8**

### Naive Samples

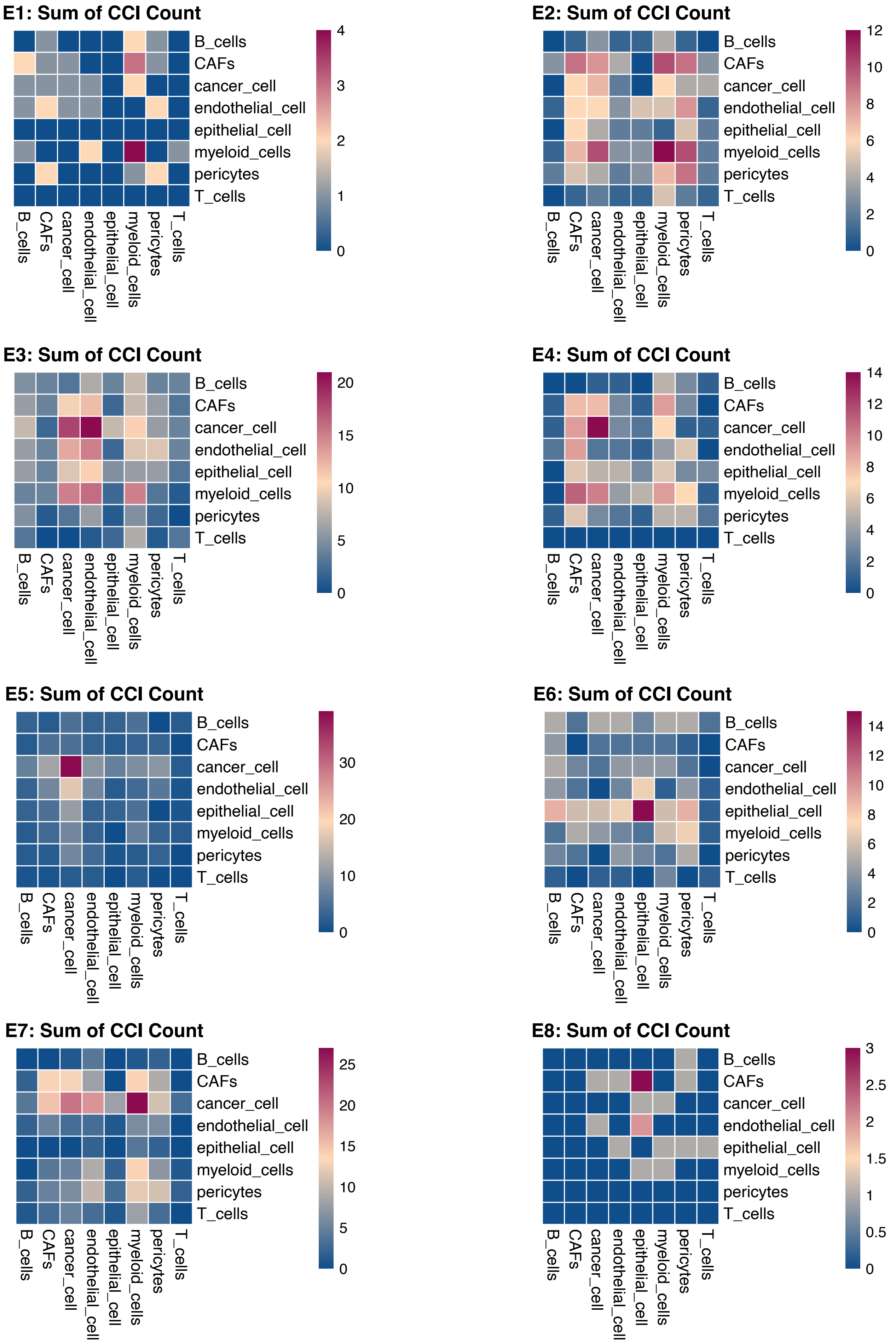

**Supplementary Figure 8**. Global cell–cell interaction (CCI) profiles for each sample within the Naïve group from the spatial transcriptomics dataset. Each heatmap corresponds to a single sample (E1, E2, E3, E4, E5, E6, E7, E8), with rows representing ligand-expressing (sender) cell types and columns representing receptor-expressing (receiver) cell types. The color intensity reflects the total number of significant ligand–receptor interactions detected between each cell-type pair.

**Supplementary Figure 9**

### SBRT Samples

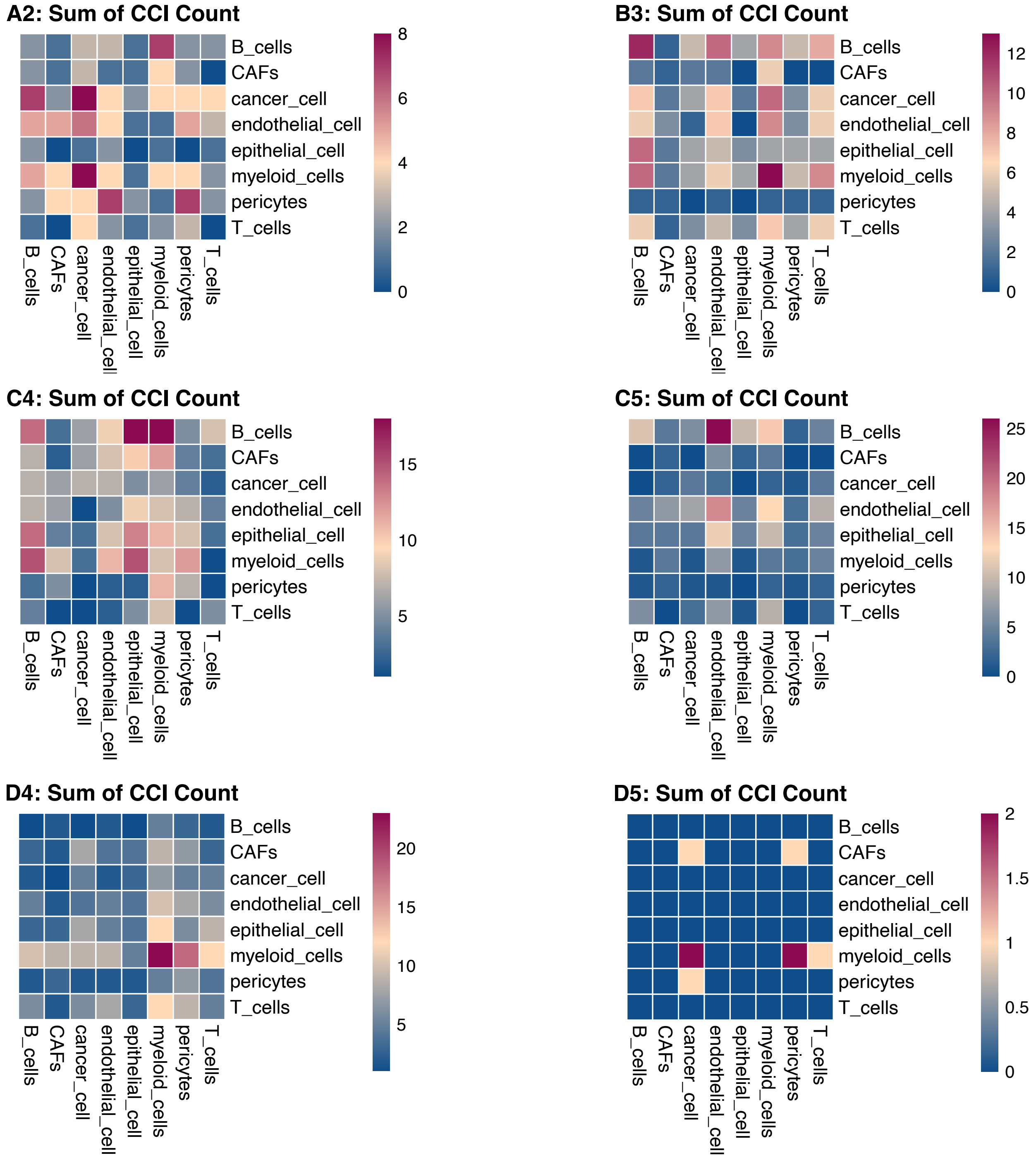


**Supplementary Figure 9**. Global cell–cell interaction (CCI) profiles for each sample within the SBRT treatment group from the spatial transcriptomics dataset. Each heatmap corresponds to a single sample (A2, B3, C4, C5, D4, D5), with rows representing ligand-expressing (sender) cell types and columns representing receptor-expressing (receiver) cell types. The color intensity reflects the total number of significant ligand–receptor interactions detected between each cell-type pair.

**Supplementary Figure 10**

### CRT30 Samples

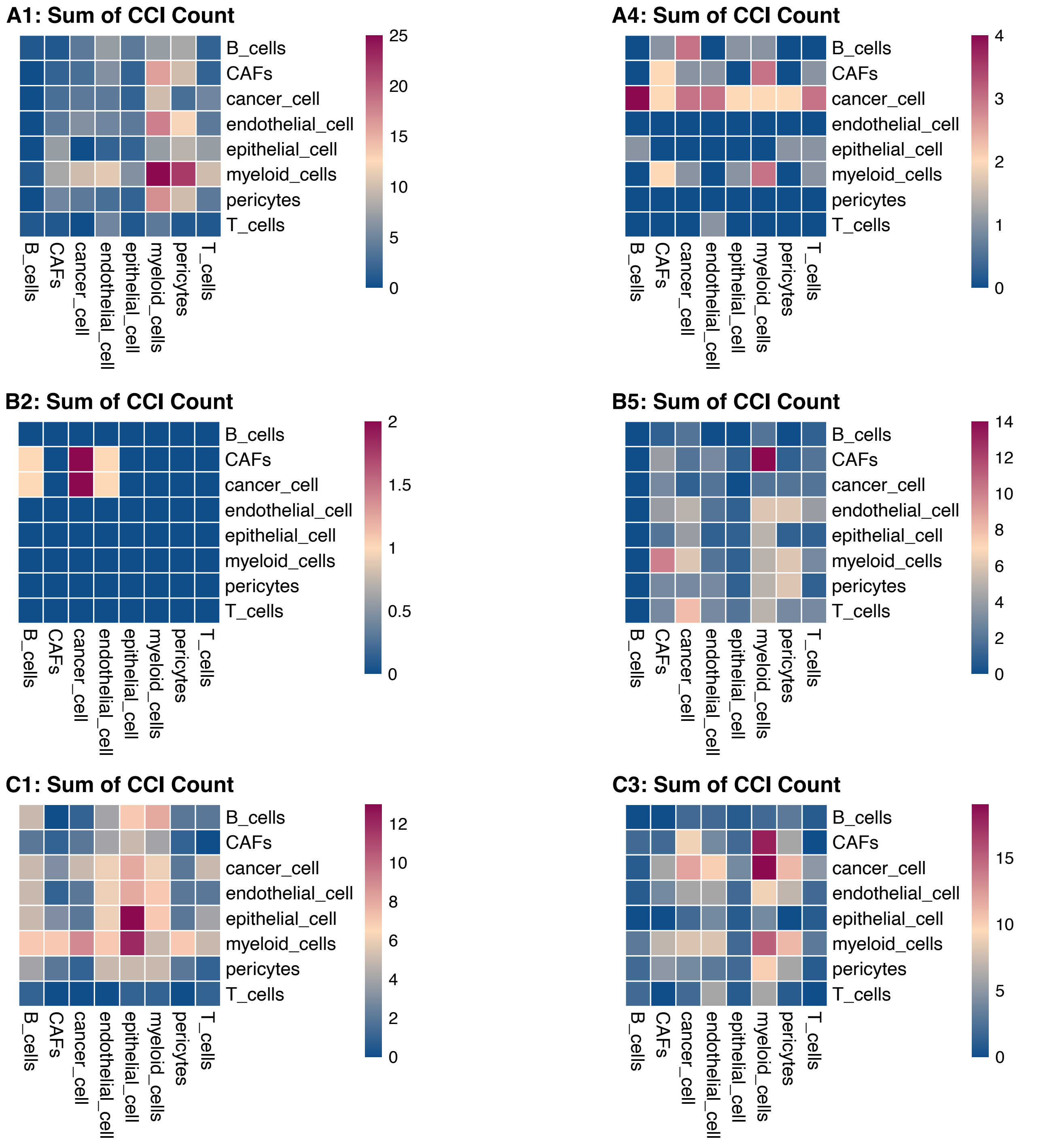


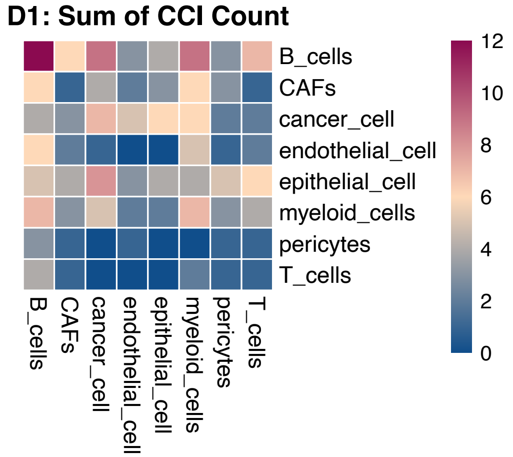


**Supplementary Figure 10.** Global cell–cell interaction (CCI) profiles for each sample within the CRT30 treatment group from the spatial transcriptomics dataset. Each heatmap corresponds to a single sample (A1, A4, B2, B5, C1, C3, D1), with rows representing ligand-expressing (sender) cell types and columns representing receptor-expressing (receiver) cell types. The color intensity reflects the total number of significant ligand–receptor interactions detected between each cell-type pair.

**Supplementary Figure 11**

### CRT50 Samples

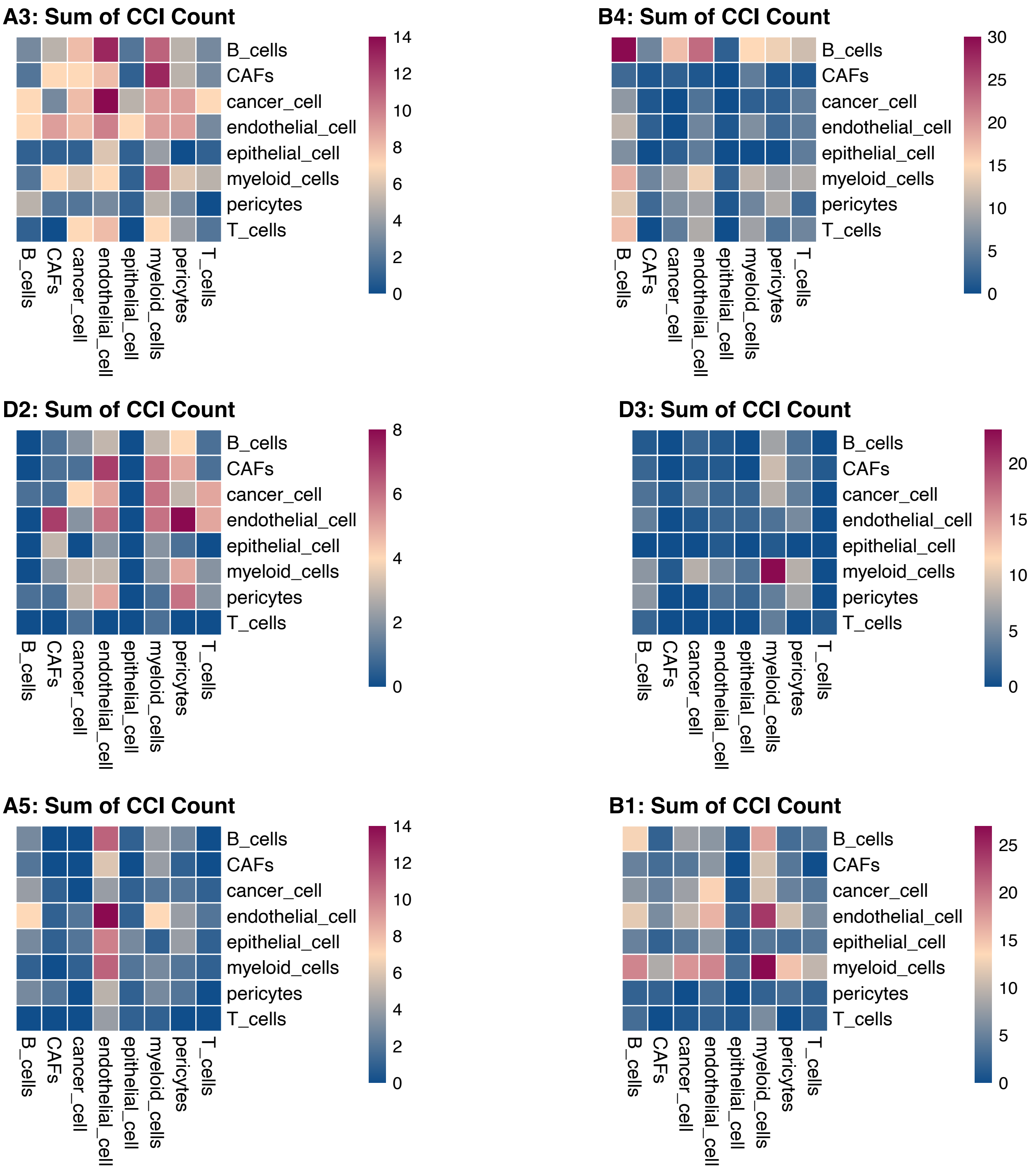


**Supplementary Figure 11**. Global cell–cell interaction (CCI) profiles for each sample within the CRT50 treatment group from the spatial transcriptomics dataset. Each heatmap corresponds to a single sample (A3, B4, D2, D3, A5, B1), with rows representing ligand-expressing (sender) cell types and columns representing receptor-expressing (receiver) cell types. The color intensity reflects the total number of significant ligand–receptor interactions detected between each cell-type pair.

#### Supplementary Figure 12

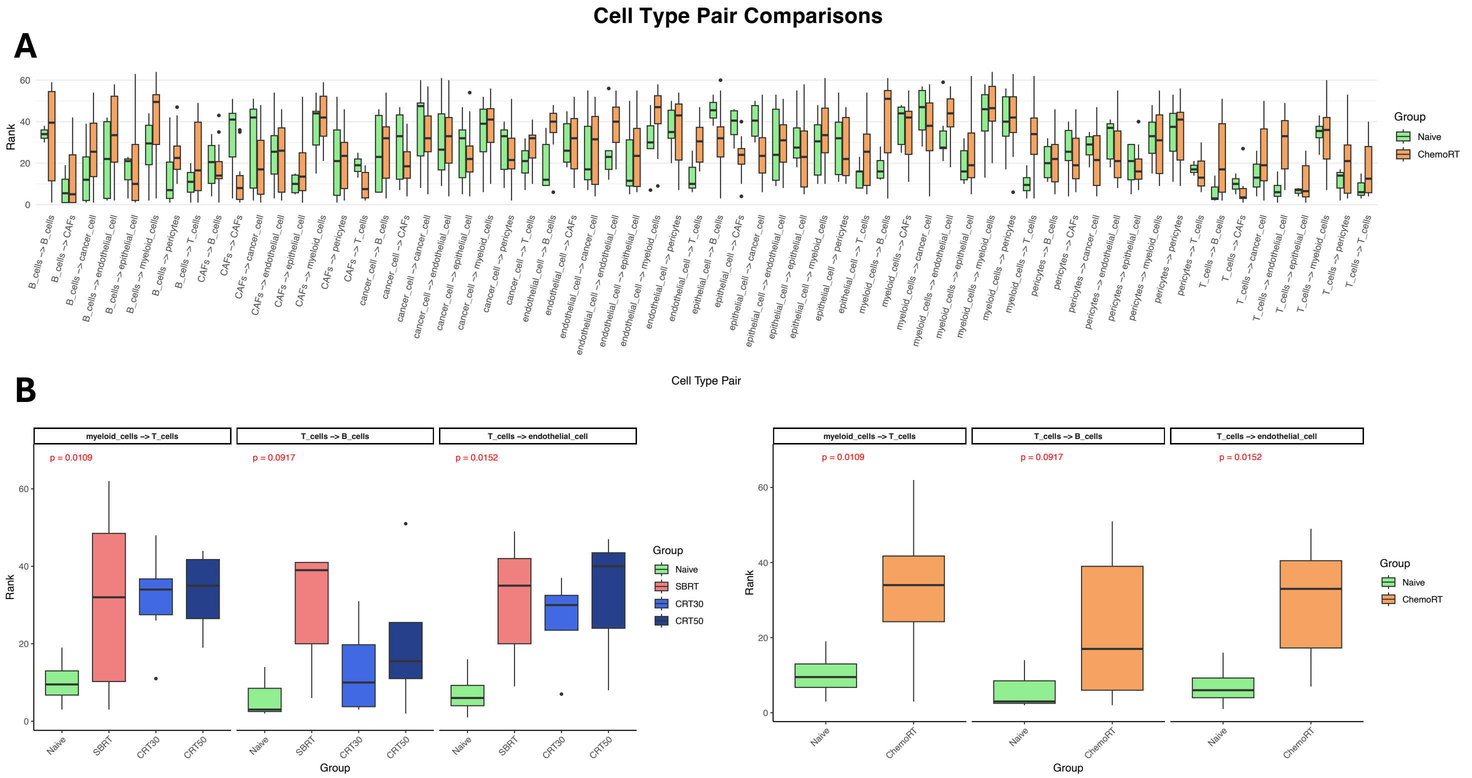


**Supplementary Figure 12**. (A) Box plots showing the distribution of interaction ranks for each cell type pair across all samples in the Naive and ChemoRT groups, based on the spatial transcriptomics dataset. Ranks were computed within each sample to reflect the relative abundance of ligand–receptor interactions. Only cell-type pairs present across all treatment groups were retained for comparison. (B) Statistical comparison of selected cell type pairs across treatment conditions. Interaction ranks were compared using the Mann–Whitney U test (Wilcoxon rank-sum test), with p-values shown in red.

#### Supplementary Figure 13

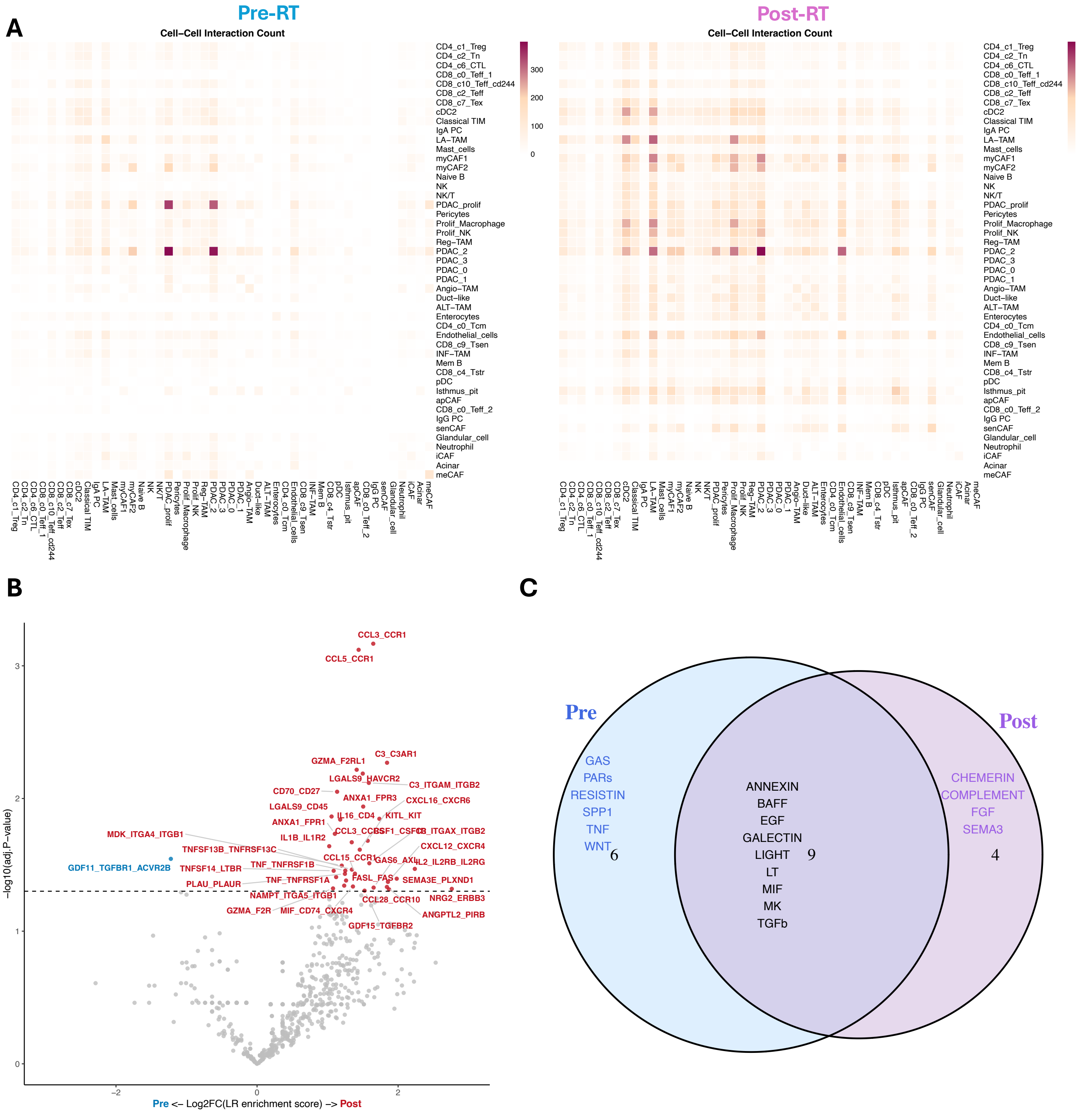


**Supplementary Figure 13**. (A) Heatmaps showing the global cell–cell interaction (CCI) counts from the single-cell RNA-seq dataset, comparing pre-RT and post-RT conditions. In each heatmap, rows represent ligand-expressing (sender) cell types, and columns represent receptor-expressing (receiver) cell types. Color intensity reflects the number of significant ligand–receptor interactions detected between each pair. (B) Volcano plot illustrating differentially enriched ligand–receptor (L–R) interactions between pre-RT and post-RT groups. Red-labeled interactions represent those significantly enriched post-RT, while blue-labeled interactions are enriched pre-RT. (C) Venn diagram showing pathway-level enrichment of CCIs involving the PDAC_3 cell type in the single-cell RNA-seq data. Pathways uniquely enriched pre-RT (blue), post-RT (purple), and those shared between both conditions (overlap) are indicated. Post-RT–enriched pathways include SEMA3, COMPLEMENT, and CHEMERIN, while pre-RT–specific pathways include GAS, SPP1, and WNT.

### Supplementary Figure 14

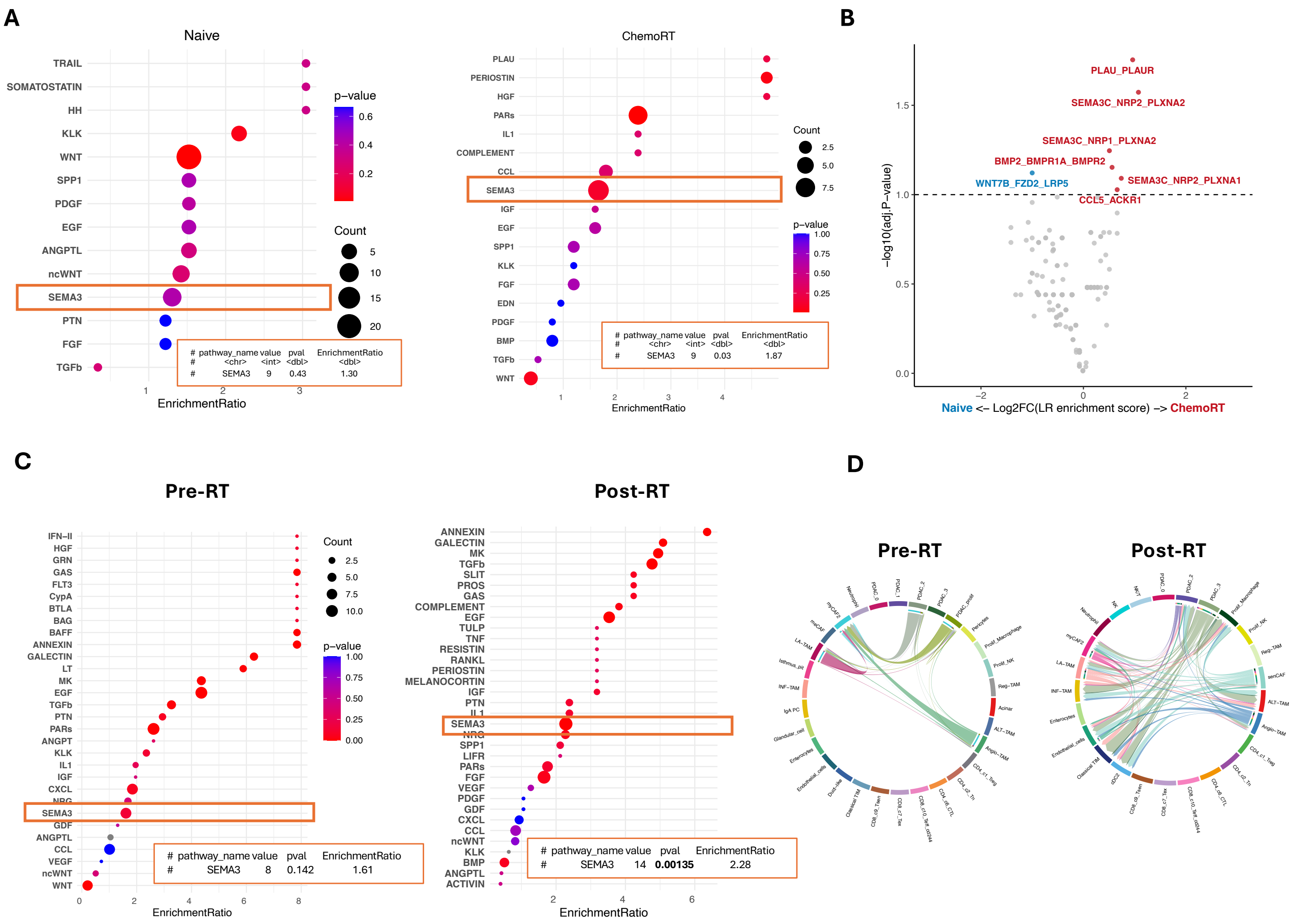


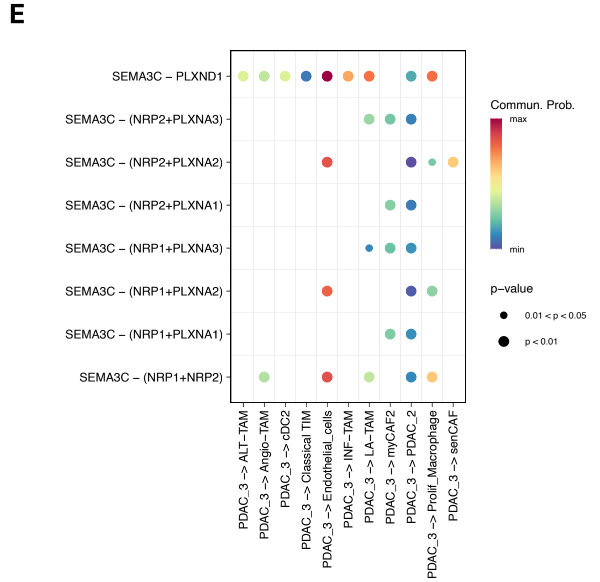


**Supplementary Figure 14**. (A) Dot plots showing enrichment of ligand–receptor (L–R) signaling pathways from spatial transcriptomics data. Notably, SEMA3 signaling is elevated in ChemoRT samples (enrichment ratio (ER): 1.30, p = 0.03) compared to treatment-naïve (ER: 1.87, p = 0.43). (B) Volcano plot of L-R interactions in PDAC_3 persister cells highlighting that the most significantly upregulated interactions post-ChemoRT are dominated by SEMA3-related pairs. (C) Dot plots from single-cell RNA-seq analysis provide orthogonal validation, showing increased SEMA3 pathway activity after RT (ER: 2.28, p = 0.00135) compared to pre-RT (ER: 1.61, p = 0.142). (D) Circos plots illustrating specific SEMA3 interactions from scRNA-seq data, particularly between PDAC_3 cells and fibroblast subtypes (senCAF, myCAF2) and myeloid subtypes (ALT-TAMs). (E) Dot plot detailing the specific cell-type–level interactions involving SEMA3 ligand-receptor pairs, with communication probabilities and significance levels indicated.

#### Supplementary Figure 15

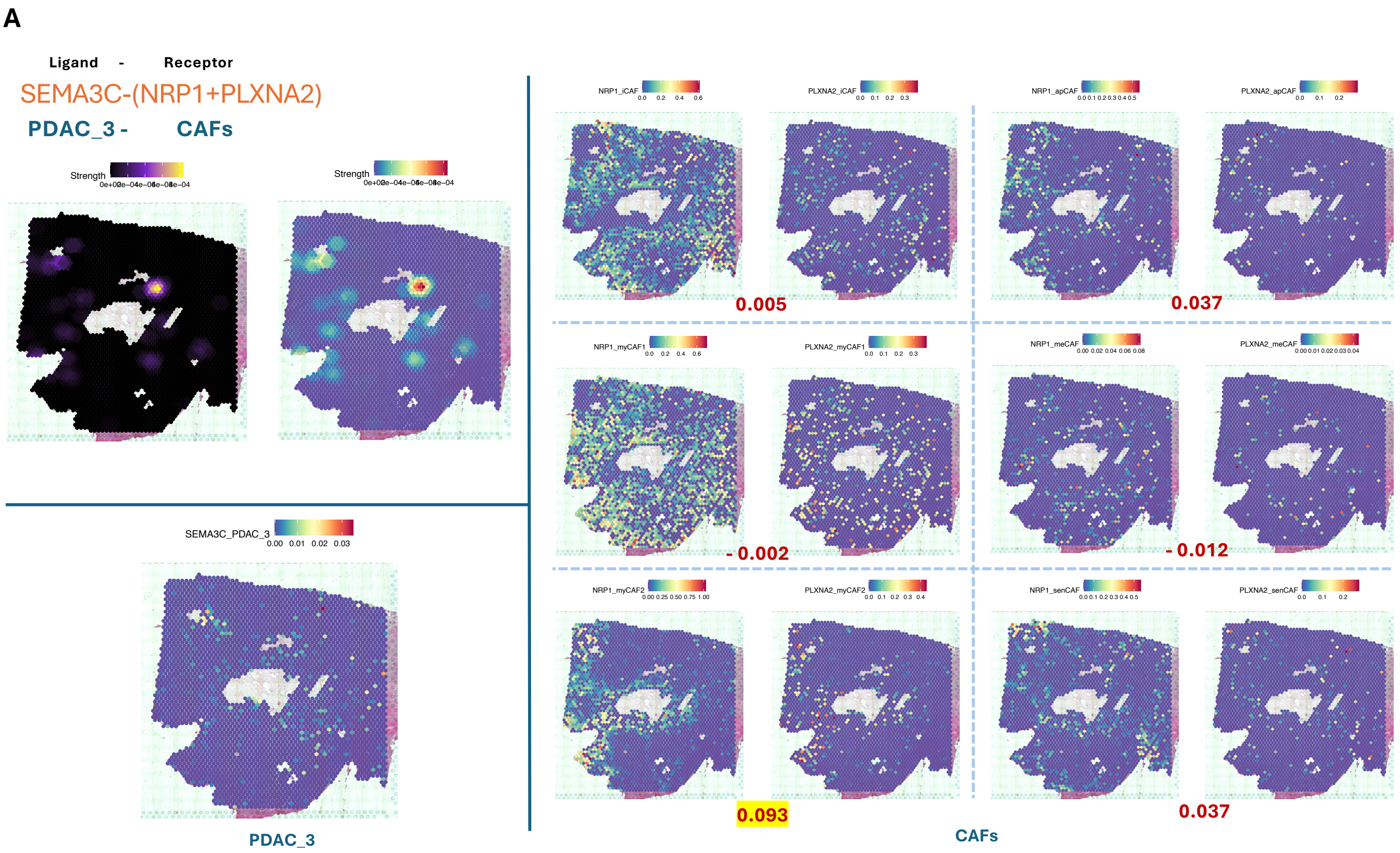


**B**

Regression on CCI strength with NRP1 expression

```
lm(formula = Strength ~ iCAF + apCAF + meCAF + myCAF1 + myCAF2 +
    senCAF, data = data)

Residuals:
       Min         1Q     Median         3Q        Max
-6.440e-05 -1.812e-05 -1.812e-05 -1.148e-05  8.183e-04

Coefficients:
              Estimate Std. Error t value Pr(>|t|)
(Intercept)  1.812e-05  1.398e-06  12.960  < 2e-16 ***
iCAF        -3.690e-06  2.072e-05  -0.178   0.8587
apCAF        5.155e-05  2.415e-05   2.135   0.0329 *
meCAF        2.356e-05  1.679e-04   0.140   0.8884
myCAF1      -2.358e-05  1.290e-05  -1.827   0.0678 .
myCAF2       4.552e-05  1.104e-05   4.123 3.84e-05 ***
senCAF       2.080e-06  2.702e-05   0.077   0.9387
---
Signif. codes:  0 ‘***’ 0.001 ‘**’ 0.01 ‘*’ 0.05 ‘.’ 0.1 ‘ ’ 1

Residual standard error: 6.086e-05 on 3274 degrees of freedom
Multiple R-squared:  0.00898,   Adjusted R-squared:  0.007163
F-statistic: 4.944 on 6 and 3274 DF,  p-value: 4.786e-05
```

Regression on CCI strength with PLXNA2 expression

```
lm(formula = Strength ~ iCAF + apCAF + meCAF + myCAF1 + myCAF2 +
    senCAF, data = data)

Residuals:
       Min         1Q     Median         3Q        Max
-1.521e-04 -1.744e-05 -1.744e-05 -1.406e-05  8.274e-04

Coefficients:
              Estimate Std. Error t value Pr(>|t|)
(Intercept)  1.744e-05  1.122e-06  15.542  < 2e-16 ***
iCAF        -1.234e-05  6.743e-05  -0.183   0.8548
apCAF        3.994e-04  7.227e-05   5.527 3.51e-08 ***
meCAF       -5.130e-04  4.580e-04  -1.120   0.2627
myCAF1      -3.032e-05  3.514e-05  -0.863   0.3884
myCAF2       6.360e-05  2.977e-05   2.137   0.0327 *
senCAF       1.731e-04  8.842e-05   1.958   0.0503 .
---
Signif. codes:  0 ‘***’ 0.001 ‘**’ 0.01 ‘*’ 0.05 ‘.’ 0.1 ‘ ’ 1

Residual standard error: 6.056e-05 on 3274 degrees of freedom
Multiple R-squared:  0.01895,   Adjusted R-squared:  0.01715
F-statistic: 10.54 on 6 and 3274 DF,  p-value: 1.286e-11
```

Regression on CCI strength with (NRP1+PLXNA2) expression

```
lm(formula = Strength ~ iCAF + apCAF + meCAF + myCAF1 + myCAF2 +
    senCAF, data = data)

Residuals:
       Min         1Q     Median         3Q        Max
-6.424e-05 -1.686e-05 -1.686e-05 -1.170e-05  8.188e-04

Coefficients:
              Estimate Std. Error t value Pr(>|t|)
(Intercept)  1.686e-05  1.436e-06  11.743  < 2e-16 ***
iCAF        -3.080e-06  1.864e-05  -0.165 0.868752
apCAF        7.639e-05  2.145e-05   3.562 0.000374 ***
meCAF       -2.108e-05  1.494e-04  -0.141 0.887811
myCAF1      -1.797e-05  1.134e-05  -1.584 0.113208
myCAF2       4.092e-05  9.449e-06   4.331 1.53e-05 ***
senCAF       1.768e-05  2.430e-05   0.728 0.466927
---
Signif. codes:  0 ‘***’ 0.001 ‘**’ 0.01 ‘*’ 0.05 ‘.’ 0.1 ‘ ’ 1

Residual standard error: 6.071e-05 on 3274 degrees of freedom
Multiple R-squared:  0.01387,   Adjusted R-squared:  0.01206
F-statistic: 7.672 on 6 and 3274 DF,  p-value: 3.332e-08
```

**Supplementary Figure 15.** (A) Spatial visualization of the localized cell-cell interaction (CCI) strength between PDAC_3 persister cells and cancer-associated fibroblasts (CAFs) mediated by the ligand-receptor pair SEMA3C–(NRP1+PLXNA2). The left panel shows the spatial distribution of interaction strength, while the middle and right panels display receptor expression (NRP1 and PLXNA2) in different CAF subtypes. Spearman correlation coefficients between receptor expression and interaction strength are shown in red.(B) We use regression models to capture the contributions of sub-cell types to cell-cell interactions. Linear regression models assess the contribution of each CAF subtype to the observed CCI strength. Three models were fitted with the interaction strength as the response and expression levels of NRP1, PLXNA2, or the combined (NRP1 + PLXNA2) as predictors, along with indicator variables for each CAF subtype. The results suggest differential contributions of CAF subtypes to ligand-receptor mediated interactions, with regression coefficients and significance values summarized in the bottom panels.

#### Supplementary Figure 16

A

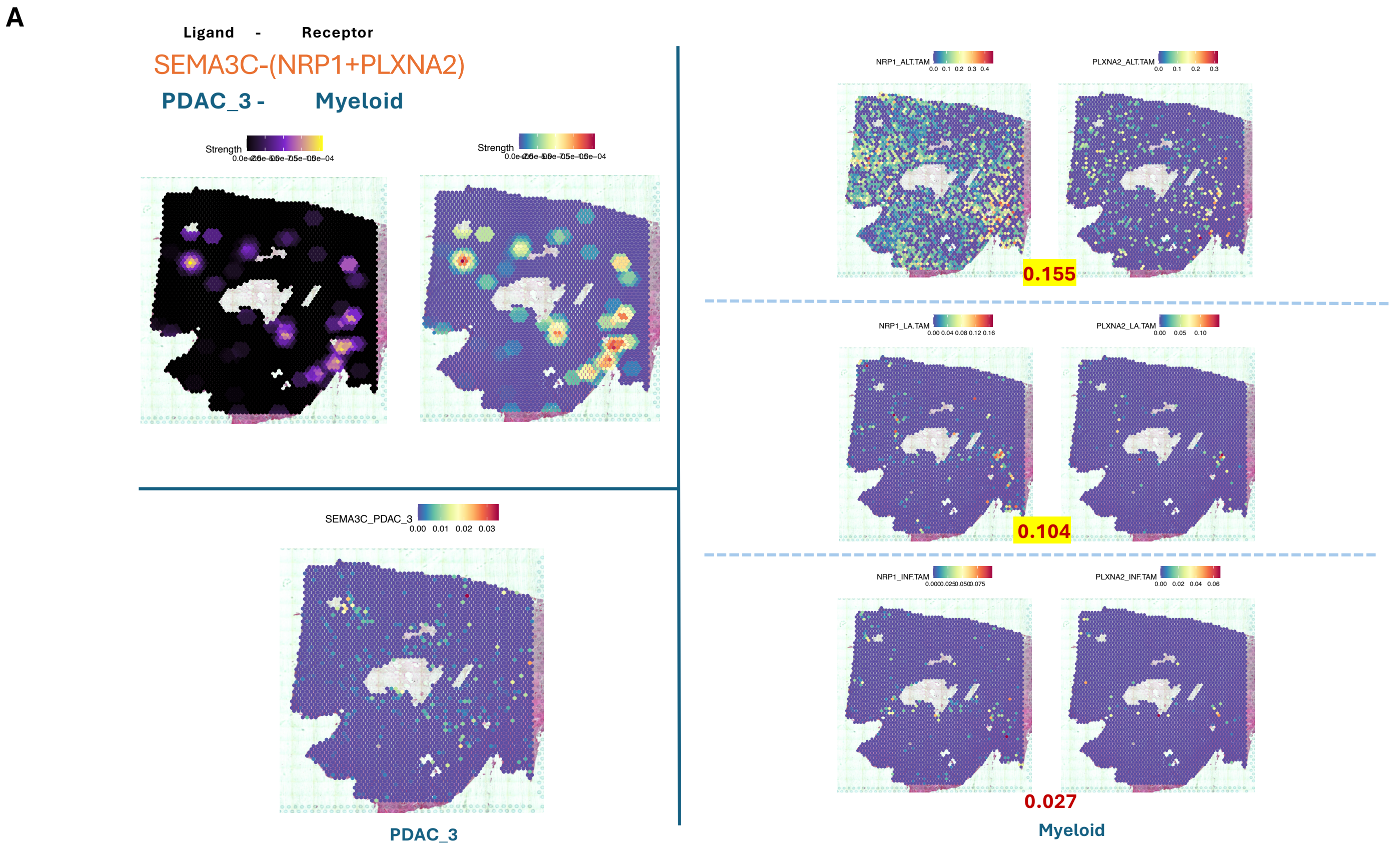


B

Regression on CCI strength with NRP1 expression

```
lm(formula = Strength ~ ALT.TAM + LA.TAM + INF.TAM, data = data)

Residuals:
       Min         1Q     Median         3Q        Max
-2.150e-05 -5.990e-06 -4.334e-06 -4.334e-06  9.133e-05

Coefficients:
              Estimate Std. Error t value Pr(>|t|)
(Intercept)  4.334e-06  3.116e-07  13.907  < 2e-16 ***
ALT.TAM      2.387e-05  3.515e-06   6.791 1.31e-11 ***
LA.TAM       6.493e-05  2.164e-05   3.001  0.00271 **
INF.TAM     -6.670e-05  5.157e-05  -1.293  0.19597
---
Signif. codes:  0 ‘***’ 0.001 ‘**’ 0.01 ‘*’ 0.05 ‘.’ 0.1 ‘ ’ 1

Residual standard error: 1.416e-05 on 3277 degrees of freedom
Multiple R-squared:  0.02006,	Adjusted R-squared:  0.01916
F-statistic: 22.36 on 3 and 3277 DF,  p-value: 2.517e-14
```

Regression on CCI strength with PLXNA2 expression

```
lm(formula = Strength ~ ALT.TAM + LA.TAM + INF.TAM, data = data)

Residuals:
       Min         1Q     Median         3Q        Max
-2.407e-05 -5.318e-06 -5.318e-06 -5.318e-06  9.167e-05

Coefficients:
             Estimate Std. Error t value Pr(>|t|)
(Intercept) 5.318e-06  2.603e-07  20.432  < 2e-16 ***
ALT.TAM     3.098e-05  8.184e-06   3.785 0.000157 ***
LA.TAM      1.880e-04  4.595e-05   4.091 4.39e-05 ***
INF.TAM     5.117e-05  1.023e-04   0.500 0.617055
---
Signif. codes:  0 ‘***’ 0.001 ‘**’ 0.01 ‘*’ 0.05 ‘.’ 0.1 ‘ ’ 1

Residual standard error: 1.419e-05 on 3277 degrees of freedom
Multiple R-squared:  0.01516,	Adjusted R-squared:  0.01426
F-statistic: 16.82 on 3 and 3277 DF,  p-value: 7.673e-11
```

Regression on CCI strength with (NRP1+PLXNA2) expression

```
lm(formula = Strength ~ ALT.TAM + LA.TAM + INF.TAM, data = data)

Residuals:
       Min         1Q     Median         3Q        Max
-2.539e-05 -6.040e-06 -3.993e-06 -3.993e-06  8.690e-05

Coefficients:
              Estimate Std. Error t value Pr(>|t|)
(Intercept)  3.993e-06  3.165e-07  12.618  < 2e-16 ***
ALT.TAM      2.416e-05  3.108e-06   7.773 1.02e-14 ***
LA.TAM       6.805e-05  1.737e-05   3.919 9.09e-05 ***
INF.TAM     -2.486e-05  4.151e-05  -0.599    0.549
---
Signif. codes:  0 ‘***’ 0.001 ‘**’ 0.01 ‘*’ 0.05 ‘.’ 0.1 ‘ ’ 1

Residual standard error: 1.409e-05 on 3277 degrees of freedom
Multiple R-squared:  0.02869,	Adjusted R-squared:  0.0278
F-statistic: 32.27 on 3 and 3277 DF,  p-value: < 2.2e-16
```

**Supplementary Figure 16.** (A) Spatial visualization of the localized cell-cell interaction (CCI) strength between PDAC_3 persister cells and myeloid cells mediated by the ligand-receptor pair SEMA3C–(NRP1+PLXNA2). The left panel shows the spatial distribution of interaction strength across the tissue section. The right panels display receptor expression (NRP1 and PLXNA2) across distinct myeloid subtypes, including ALT.TAM, LA.TAM, and INF.TAM. Spearman correlation coefficients between receptor expression and CCI strength are shown in red. (B) Linear regression models quantify the contributions of individual myeloid subtypes to the CCI strength. Three models were fitted using expression levels of NRP1, PLXNA2, or the combined (NRP1 + PLXNA2) as predictors, along with indicator variables for myeloid subtypes.

#### Supplementary Figure 17

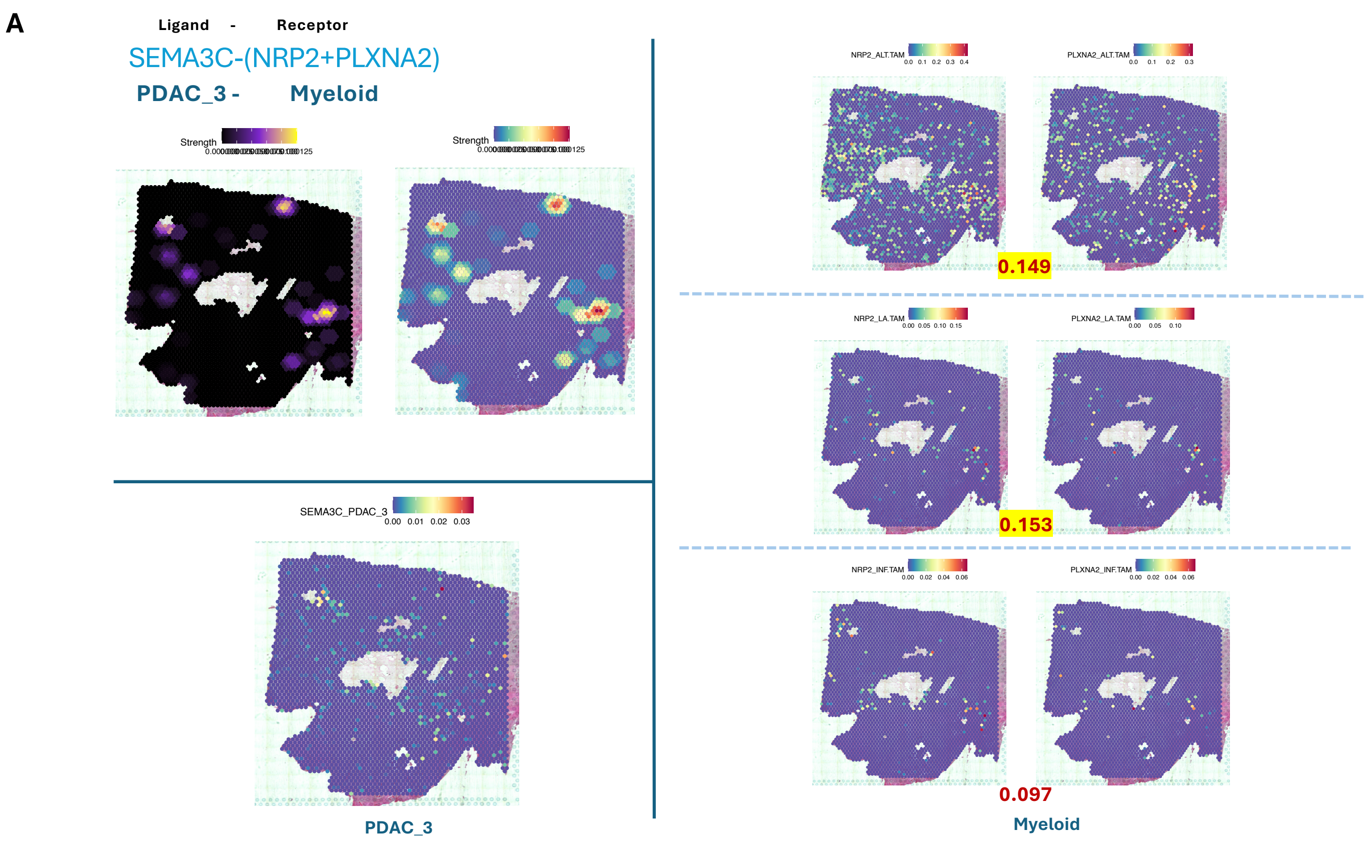


**B**

Regression on CCI strength with NRP2 expression

```
lm(formula = Strength ~ ALT.TAM + LA.TAM + INF.TAM, data = data)

Residuals:
       Min         1Q     Median         3Q        Max
-4.757e-05 -3.924e-06 -3.924e-06 -3.924e-06  1.220e-04

Coefficients:
             Estimate Std. Error t value Pr(>|t|)
(Intercept) 3.924e-06  2.635e-07  14.892  < 2e-16 ***
ALT.TAM     2.444e-05  5.397e-06   4.528 6.16e-06 ***
LA.TAM      1.499e-04  2.951e-05   5.080 3.98e-07 ***
INF.TAM     1.078e-04  6.779e-05   1.591    0.112
---
Signif. codes:  0 '***' 0.001 '**' 0.01 '*' 0.05 '.' 0.1 ' ' 1

Residual standard error: 1.377e-05 on 3277 degrees of freedom
Multiple R-squared:  0.02639,	Adjusted R-squared:  0.0255
F-statistic: 29.61 on 3 and 3277 DF,  p-value: < 2.2e-16
```

Regression on CCI strength with PLXNA2 expression

```
lm(formula = Strength ~ ALT.TAM + LA.TAM + INF.TAM, data = data)

Residuals:
       Min         1Q     Median         3Q        Max
-4.095e-05 -4.286e-06 -4.286e-06 -4.286e-06  1.216e-04

Coefficients:
             Estimate Std. Error t value Pr(>|t|)
(Intercept) 4.287e-06  2.533e-07  16.924  < 2e-16 ***
ALT.TAM     2.038e-05  7.964e-06   2.559   0.0106 *
LA.TAM      2.665e-04  4.472e-05   5.960 2.79e-09 ***
INF.TAM     1.460e-04  9.958e-05   1.466   0.1427
---
Signif. codes:  0 '***' 0.001 '**' 0.01 '*' 0.05 '.' 0.1 ' ' 1

Residual standard error: 1.381e-05 on 3277 degrees of freedom
Multiple R-squared:  0.02077,	Adjusted R-squared:  0.01987
F-statistic: 23.16 on 3 and 3277 DF,  p-value: 7.801e-15
```

Regression on CCI strength with (NRP2+PLXNA2) expression

```
lm(formula = Strength ~ ALT.TAM + LA.TAM + INF.TAM, data = data)

Residuals:
       Min         1Q     Median         3Q        Max
-4.126e-05 -3.677e-06 -3.677e-06 -3.677e-06  1.222e-04

Coefficients:
             Estimate Std. Error t value Pr(>|t|)
(Intercept) 3.677e-06  2.702e-07  13.609  < 2e-16 ***
ALT.TAM     2.245e-05  4.218e-06   5.322 1.10e-07 ***
LA.TAM      1.156e-04  1.991e-05   5.804 7.11e-09 ***
INF.TAM     1.197e-04  5.030e-05   2.380   0.0174 *
---
Signif. codes:  0 '***' 0.001 '**' 0.01 '*' 0.05 '.' 0.1 ' ' 1

Residual standard error: 1.371e-05 on 3277 degrees of freedom
Multiple R-squared:  0.03535,	Adjusted R-squared:  0.03447
F-statistic: 40.03 on 3 and 3277 DF,  p-value: < 2.2e-16
```

**Supplementary Figure 17.** (A) Spatial visualization of the localized cell-cell interaction (CCI) strength between PDAC_3 persister cells and myeloid cells, mediated by the ligand-receptor pair SEMA3C–(NRP2+PLXNA2). The left panel shows the spatial distribution of interaction strength across the tissue section. The right panels illustrate the expression levels of receptors NRP2 and PLXNA2 across myeloid subtypes ALT.TAM, LA.TAM, and INF.TAM, with corresponding Spearman correlation coefficients with CCI strength displayed in red. (B) Linear regression models evaluate the contribution of each myeloid subtype to the observed CCI strength. Three models were constructed using the expression levels of NRP2, PLXNA2, or their combination (NRP2 + PLXNA2) as predictors, alongside subtype indicators. Model outputs reveal differential involvement of myeloid subpopulations in mediating SEMA3C-driven signaling, as reflected in the regression coefficients and significance levels.
